# Inter-species geographic signatures for tracing horizontal gene transfer and long-term persistence of carbapenem resistance

**DOI:** 10.1101/2021.12.08.471225

**Authors:** Rauf Salamzade, Abigail L. Manson, Bruce J. Walker, Thea Brennan-Krohn, Colin J. Worby, Peijun Ma, Lorrie L. He, Terrance P. Shea, James Qu, Sinéad B. Chapman, Whitney Howe, Sarah K. Young, Jenna I. Wurster, Mary L. Delaney, Sanjat Kanjilal, Andrew B. Onderdonk, Alejandro Pironti, Cassiana E. Bittencourt, Gabrielle M. Gussin, Diane Kim, Ellena M. Peterson, Mary Jane Ferraro, David C. Hooper, Erica S. Shenoy, Christina A. Cuomo, Deborah T. Hung, Lisa A. Cosimi, Susan S. Huang, James E. Kirby, Virginia M. Pierce, Roby P. Bhattacharyya, Ashlee M. Earl

**Author notes:** Microbiology Doctoral Training Program, University of Wisconsin-Madison, Madison, WI, 53706.

## Abstract

**Background:** Carbapenem-resistant *Enterobacterales* (CRE) are an urgent global health threat. Inferring the dynamics of local CRE dissemination is currently limited by our inability to confidently trace the spread of resistance determinants to unrelated bacterial hosts. Whole genome sequence comparison is useful for identifying CRE clonal transmission and outbreaks, but high-frequency horizontal gene transfer (HGT) of carbapenem resistance genes and subsequent genome rearrangement complicate tracing the local persistence and mobilization of these genes across organisms.

**Methods:** To overcome this limitation, we developed a new approach to identify recent HGT of large, near-identical plasmid segments across species boundaries, which also allowed us to overcome technical challenges with genome assembly. We applied this to complete and near-complete genome assemblies to examine the local spread of CRE in a systematic, prospective collection of all CRE, as well as time- and species-matched carbapenem susceptible *Enterobacterales*, isolated from patients from four U.S. hospitals over nearly five years.

**Results:** Our CRE collection comprised a diverse range of species, lineages and carbapenem resistance mechanisms, many of which were encoded on a variety of promiscuous plasmid types. We found and quantified rearrangement, persistence, and repeated transfer of plasmid segments, including those harboring carbapenemases, between organisms over multiple years. Some plasmid segments were found to be strongly associated with specific locales, thus representing *geographic signatures* that make it possible to trace recent and localized HGT events.

Functional analysis of these signatures revealed genes commonly found in plasmids of nosocomial pathogens, such as functions required for plasmid retention and spread, as well survival against a variety of antibiotic and antiseptics common to the hospital environment.

**Conclusions:** Collectively, the framework we developed provides a clearer, high resolution picture of the epidemiology of antibiotic resistance importation, spread, and persistence in patients and healthcare networks.

## Background

Carbapenem-resistant *Enterobacterales* (CRE) cause difficult-to-treat infections [1–5] with high mortality rates [6–8], largely because antibiotic options for treating them are limited [9, 10]. CRE are also highly transmissible through contact [11–16], leading to nosocomial outbreaks that are costly to contain with significant patient morbidity and mortality [17–19], making CRE a leading healthcare problem [20–25]. Despite the adoption of extensive infection control measures [11, 12] that have begun to curb the incidence of CRE infection in some countries, the global incidence of CRE infections continues to rise [20,23,24,26–28].

Genomic studies are providing new insights into the emergence and spread of CRE within healthcare institutions [29–31]. Comprising many species, most commonly *Klebsiella pneumoniae*, *Escherichia coli*, and *Enterobacter cloacae* complex, CRE can be found in diverse environments within hospitals [32–35], ranging from the gastrointestinal tract of asymptomatic carriers [36, 37] to contaminated hospital sinks and drains [38–42]. CRE are readily acquired and spread from these reservoirs [13, 40], including hospital-adapted *high-risk lineages* (*e.g. K. pneumoniae* sequence type (ST)-258) [43] that are associated with both intra- and inter-facility clonal transmission [13,29,44,45]. Known reservoirs tend to be polymicrobial and thus can act as sites for CRE diversification and are believed to have played important roles in horizontal gene transfer (HGT) of carbapenemases [40, 42].

Though bioinformatic approaches targeting the sequences surrounding carbapenemases have been used to predict the movement of carbapenemases across some CRE populations [46], tracing the movement and persistence of these genes within facilities is complicated. CRE reservoirs can be large and diverse, and comparatively few have been studied [39,40,42,47,48]. Furthermore, HGT rates are predicted to be high [49–51], and carbapenemase containing plasmids frequently recombine at sites of repetitive sequence, leading to mosaic plasmid structures [31, 52]. Given this, carbapenemase-containing plasmid sequences are challenging to accurately assemble. Long-read sequencing technologies have provided the strongest evidence for HGT of plasmids containing carbapenemases between *Enterobacterales* within individual hospitals, as well as for transmission between patients and hospital reservoirs [40, 42]. However, it remains challenging to trace carbapenemase movement within and between different plasmid backgrounds and organisms even when using high quality assemblies generated using long-read sequencing data.

We previously published results from a surveillance study conducted in 2012-2013 [30] that highlighted the shortcomings of existing methods for tracking CRE movement. Here, we expanded this initial study to capture all patient-derived CRE, regardless of infection site or resistance mechanism, from across the same four hospitals over an additional three-year period from December 2013 through 2016. Our sequencing methodology, which combined short paired-end and long-insert mate pair Illumina sequencing libraries, enabled high quality whole genome and plasmid assemblies for over 600 isolates. We developed a novel computational methodology to holistically screen for conserved segments within mosaic plasmids that allowed us to trace the movement and persistence of genes, including carbapenemases, within facilities. This approach revealed near-identical plasmid segments, including carbapenemase encoding segments, that crossed plasmid and species boundaries. Many of these were specific to and recurrent within a single hospital site, revealing extensive linkages between patient isolates that would have been missed otherwise. From long-read sequencing of select isolates, we also observed rapid plasmid mosaicism, including the shuffling of segments into new genomic locations, occurring on the same timescale as single nucleotide variation.

## Results

### Comprehensive collection of clinical specimen CREs from microbiology labs over a nearly five year period reveals striking diversity and clusters

As part of our continued surveillance of CRE at four large tertiary hospitals located in Boston, MA and Orange, CA, we collected and sequenced the genomes of all carbapenem-resistant *Enterobacterales* (CRE; defined here as meropenem MIC ≥ 2μg/ml; Materials and Methods) cultured from clinical specimens between December 2013 and December 2016, regardless of species or resistance mechanism. These 146 CRE were added to our earlier published dataset [30] of 74 CRE prospectively collected between August 2012 and November 2013, and 47 historical CRE isolates from the same hospitals, including 12 sets of related same-patient isolates (Figure 1; Table 1; Tables S1-S3). For each CRE, we also collected and sequenced at least one species- and time-matched carbapenem susceptible *Enterobacterales* (CSE; defined here as meropenem MIC < 2μg/ml). This collection strategy gave us access to isolates representing a wide range of carbapenem resistance mechanisms, as well as a snapshot of the sympatric susceptible population at each hospital [30] (Table S1). Consistent with our previous findings, CRE were most often cultured from urine specimens (40%), followed by respiratory tract (18%), blood (9%), and bile (8%) specimens (Table S1).

**Figure 1:**
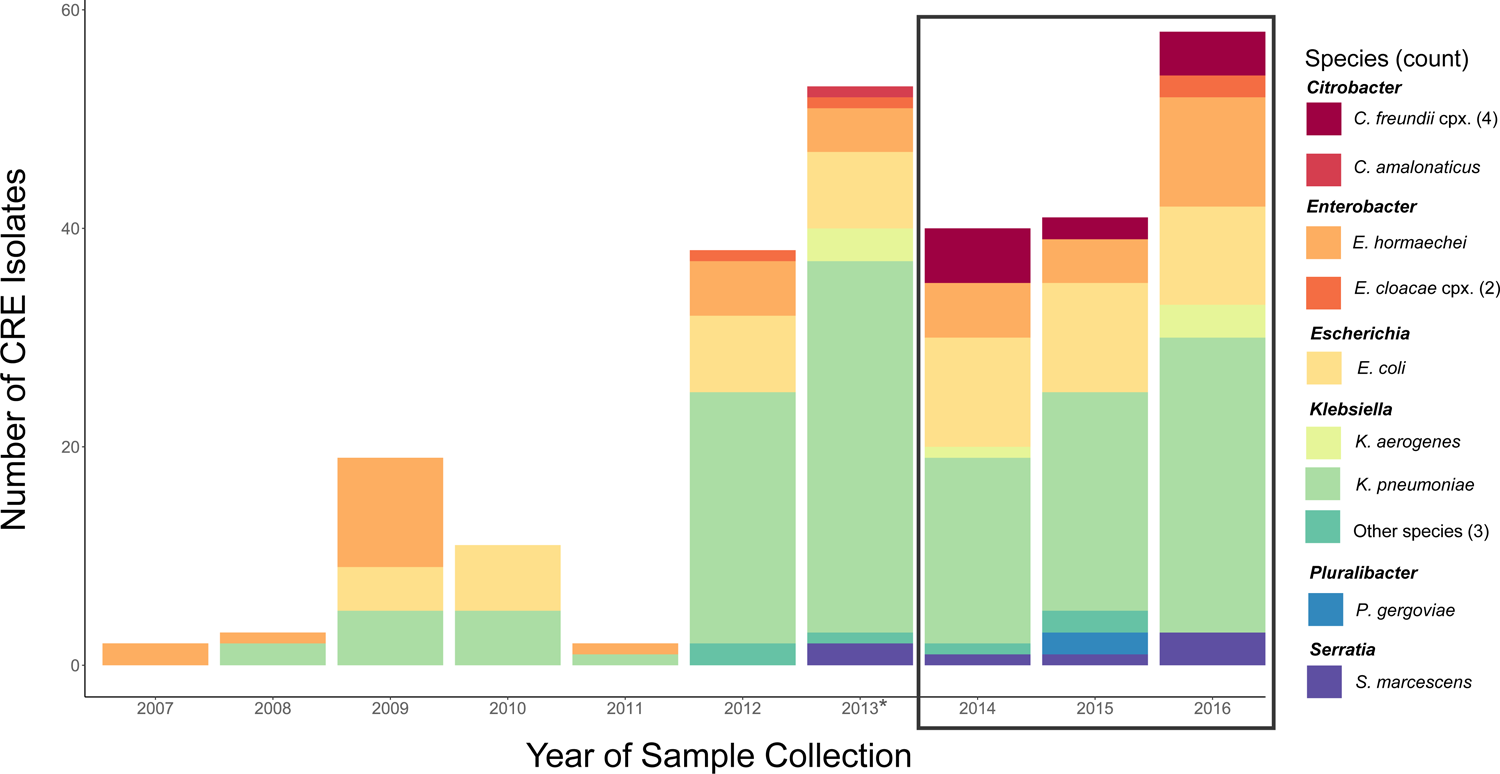
High species diversity across CRE isolates. The number of isolates collected and sequenced by year and species is shown. The black box indicates the isolates newly sequenced as part of this study, together with an additional 15 isolates from 2013 (*). All others were previously described [30].

**Table 1:** Number of isolates in the three different collections of our dataset, stratified by resistance status.

|  |  |  | Same Patient isolates |  |  |
| --- | --- | --- | --- | --- | --- |
|  | Total CRE | Total CSE | Unique patients | Total same-patient CRE isolates | Total same-patient CSE isolates |
| Boston historical collection (Cerqueira et al., 2017) | 47 | 2 | 7 | 17 | 1 |
| Initial prospective Collection (Cerqueira et al., 2017) | 74 | 138 | 5 | 7 | 3 |
| Newly sequenced prospective collection | 146 | 201 | 0 | 0 | 0 |
| Total | 267 | 341 | 12 | 24 | 4 |

For each isolate, we generated highly contiguous *de novo* genome assemblies (average scaffold N50 of 3.8 Mb; Table S4) using a combination of Illumina short paired-end and long-insert mate pair libraries [53]. Echoing our previous work [30], and consistent with previous studies [29–31,54], whole genome average nucleotide identity (ANI) comparisons revealed a vast taxonomic diversity among patient CRE isolates: 16 different species were observed among this collection, though *K. pneumoniae sensu stricto* (52%), *E. coli* (18%) and *Enterobacter hormaechei* (13%; a member of *E. cloacae* complex) were most prevalent (Figure 1; Table S1). As expected by the study design, CSE isolates were similarly distributed by taxonomy (Table S1). To further classify and explore the diversity of isolates, we generated single copy core phylogenetic trees for each species, computationally defined lineages using phylogenetic distances as previously described [31], and mapped these lineages to existing STs, including those previously defined as high-risk [43] due to their ability to cause severe and/or recurrent drug-resistant infections and rapidly spread (Materials and Methods). This revealed a striking intra-species diversity of organisms within our dataset (Figure 2; Figure S1; Table S1), though we also observed closely related clusters of isolates. Twenty percent of isolates were separated from another isolate by two or fewer single nucleotide variants (SNVs), based on core genome comparisons, which was the range of SNVs observed for related same-patient isolates (Table S3). Another 14% were separated from another isolate by a maximum of 10 core SNVs, a number previously used to suggest recent common ancestry [55, 56].

**Figure 2.**
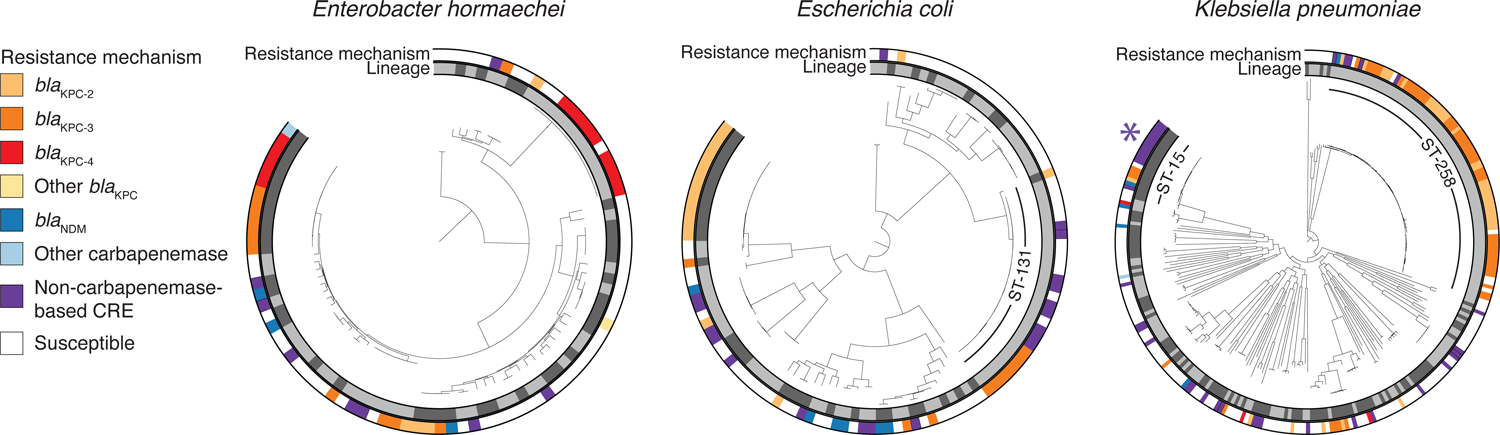
Resistance mechanisms were phylogenetically dispersed but their carriage varied among closely related isolates. For *E. hormaechei*, *E. coli*, and *K. pneumoniae*, lineages are indicated in the inner ring with alternating shades of grey. Resistance mechanisms for each isolate are shown in the outer ring. The high risk lineages *K. pneumoniae* ST-258 and *E. coli* ST-131 are marked. *K. pneumoniae* ST-15, which contained an example of likely nosocomial spread of porin-based CRE, is also marked with an asterisk.

### Heterogeneous phylogenetic distribution of resistance determinants, and evidence for nosocomial spread of non-CP-CRE

Our analysis of the mechanisms of carbapenem resistance revealed that the majority of resistance was mediated by carbapenemases (CP-CRE), though extended-spectrum beta lactamases coinciding with porin disruptions also contributed (Figure 2; Figure S1-S3; Table 2; Table S1, S5-S7; Supplementary Results). Pointing to the known role of HGT in the spread of carbapenem resistance in the *Enterobacterales* [30,42,52], species phylogenies revealed both carriage of identical carbapenemase alleles (and their transposon Tn*4401* variants; Table S8) among distantly related isolates and heterogeneity in the carriage of resistance genes among closely related isolates (Figure 2). Importantly, despite capturing a very small fraction of patient CSE from these hospitals, we observed CP-CRE and susceptible isolates separated by as few as 4 core SNVs, connections that would be missed by phenotyping alone (Table S9), and pointing to transfer of carbapenemase-containing plasmids within locally circulating populations.

**Table 2:** Resistome categories for carbapenem-resistant isolates in our prospective collection

| Resistance Mechanism | # of CRE isolates | # of CSE isolates | % of total isolates in this category that are CRE | % of CRE isolates with this mechanism |
| --- | --- | --- | --- | --- |
| **i. Carbapenemases** | **145** | **3** | **98.0%** | **65.9%** |
| **All bla<sub>KPC</sub>** | **124** | **2** | **98.4%** | **56.4%** |
| bla<sub>KPC-3</sub> | 60 | 1 | 98.4% | 27.3% |
| bla<sub>KPC-2</sub> | 50 | 1 | 98.0% | 22.7% |
| bla<sub>KPC-4</sub> | 11 | 0 | 100.0% | 5.0% |
| bla<sub>KPC</sub> (read based)<sup>a</sup> | 3 | 0 | 100.0% | 1.4% |
| **All bla<sub>NDM</sub>** | **14** | **0** | **100.0%** | **6.4%** |
| bla<sub>NDM-1</sub> | 11 | 0 | 100.0% | 5.0% |
| bla<sub>NDM-5</sub> | 3 | 0 | 100.0% | 1.4% |
| **All bla<sub>SME</sub>** | **5** | **1** | **83.3%** | **2.3%** |
| bla<sub>SME-1</sub> | 2 | 1 | 66.7% | 0.9% |
| **bla<sub>SME-2</sub>** | **3** | **0** | **100.0%** | **1.4%** |
| **Other carbapenemases** | **2** | **0** | **100.0%** | **0.9%** |
| **bla<sub>IMP-84</sub> and bla<sub>KPC-4</sub>** | **1** | **0** | **100.0%** | **0.5%** |
| **bla<sub>IMP-4</sub>** | **1** | **0** | **100.0%** | **0.5%** |
| **ii. ESBL/AmpC + porin defect(s)<sup>b</sup>** | **61** | **32** | **65.6%** | **27.7%** |
| **Includes any ESBL** | **46** | **11** | **80.7%** | **20.9%** |
| **Includes bla<sub>CTX-M-15</sub>** | **34** | **2** | **94.4%** | **15.5%** |
| **Includes bla<sub>SHV-12</sub>** | **11** | **5** | **68.8%** | **5.0%** |
| **Includes other ESBLs** | **13** | **5** | **72.2%** | **5.9%** |
| **Includes any AmpC** | **31** | **23** | **57.4%** | **14.1%** |
| **Includes bla<sub>EC</sub>** | **19** | **8** | **70.4%** | **8.6%** |
| **Includes bla<sub>CMY-2</sub>** | **12** | **4** | **75.0%** | **5.5%** |
| **Includes other AmpCs** | **5** | **12** | **29.4%** | **2.3%** |
| **iii. ESBL/AmpC without porin defect** | **6** | **168** | **3.4%** | **2.7%** |
| **iv. Porin defect(s) without ESBL/AmpC** | **8** | **23** | **25.8%** | **3.6%** |
| **Defect in both major porins** | **5** | **5** | **50.0%** | **2.3%** |
| **v. No known resistant determinants** | **0** | **113** | **0.0%** | **0.0%** |

<sup>a</sup>The *bla<sub>KPC</sub>* was not present in the whole genome assembly, but we were able to detect its presence by examining the read data (see Materials and Methods).
<sup>b</sup>Strains may have multiple predicted AmpC and/or ESBL genes. See Tables S17 and S18 for specific resistance determinants predicted for each isolate.

While the majority of closely related clusters (< 10 SNPs) comprised CP-CRE, we also found evidence for the nosocomial spread of non-CP-CRE. A cluster of 12 *K. pneumoniae* ST-15 isolates (separated by 0-2 SNVs) that all harbored the ESBL *bla*_CTX-M-15_ were isolated from 12 unique patients from a single hospital across four years. Eleven of 12 members also carried inactivating mutations in both major porins, OmpK35 and OmpK36 (Table S10). Though similar to an earlier report from hospitals in Greece [57], where 19 clonally-related isolates carried *bla*_CTX-M-15_, together with consistent disruptions (*ompK35*) or mutations (*ompK36*) in porins, we observed that, while each isolate carried the same, presumably vertically transmitted, frame-shifted copy of *ompK35*, all but one carried a distinct inactivated form of *ompK36* with non-identical IS element insertions [58–60].

### Geographically widespread, rapidly rearranging plasmid groups prevented accurate tracing of carbapenemase exchange between unrelated organisms

Because our assemblies were generated using data from both short paired-end and long-insert mate pair Illumina libraries, they were highly contiguous (*i.e.*, >20% of >2 kb plasmid scaffolds were predicted to represent complete, circular plasmids (Table S11)), which allowed us to examine the distribution of plasmids across isolates, genera, and geographic locations.

The vast majority of plasmids (83%) were assigned to one of 215 plasmid groups by MOB-suite (Figure S4; Table S12), of which nearly 20% contained instances of plasmids encoding carbapenemases. However, most (80%) of these carbapenemase-containing plasmid groups also contained plasmids lacking a carbapenemase (Figure S5, panel B), showcasing the known genetic flexibility of plasmids [31,54,61]. Furthermore, plasmid groups were also remarkably geographically widespread [62], with nearly all (92%) of the most prevalent plasmid groups found in isolates from multiple hospitals, including all of those containing carbapenemases (Figure S5, panel C), with a majority (70%) found in isolates from both cities.

The widespread geographic distribution of plasmids, and the ample opportunity they have to interact with and rearrange genetic content with other plasmids (Supplementary Results), complicates the tracing of clinically important resistance genes contained within plasmids. Specific genetic markers have previously been used to trace the local spread of *bla*_KPC_ [46]. However, when applied to our dataset, these markers, including specific plasmid groups, Tn*4401* isoforms and their 5 base pair (bp) or longer flanks, were also mostly geographically widespread (Figure S6).

### Identification of geographic signatures allows for local tracing of HGT and inter-molecular movement of genes

Inspired by analyses investigating larger flanking regions surrounding resistance determinants [63], we developed *ConSequences*, a broader, gene-agnostic approach to identify highly conserved and contiguous segments on plasmids that may serve as markers of local HGT of clinically important genes, such as those encoding carbapenemases and other hospital adaptive traits (https://github.com/broadinstitute/ConSequences). We first searched predicted plasmids from across our entire dataset of >600 isolates for 10 kb or larger segments that were conserved in gene order and nucleotide identity (≥ 99% per 10kb block) across two or more plasmid scaffolds (Figure 3, Figure S7-S8). Of the 4,605 unique segments meeting these criteria, 95% exhibited some amount of overlap or nesting with other segments, and 58% were identified in multiple plasmid groups, highlighting the frequent recombination between plasmids and complex nesting among mobile genetic elements in *Enterobacterales* [31, 64]. The size of conserved segments ranged from 10 kb to 310 kb, with longer segments typically observed in fewer isolates.

**Figure 3.**
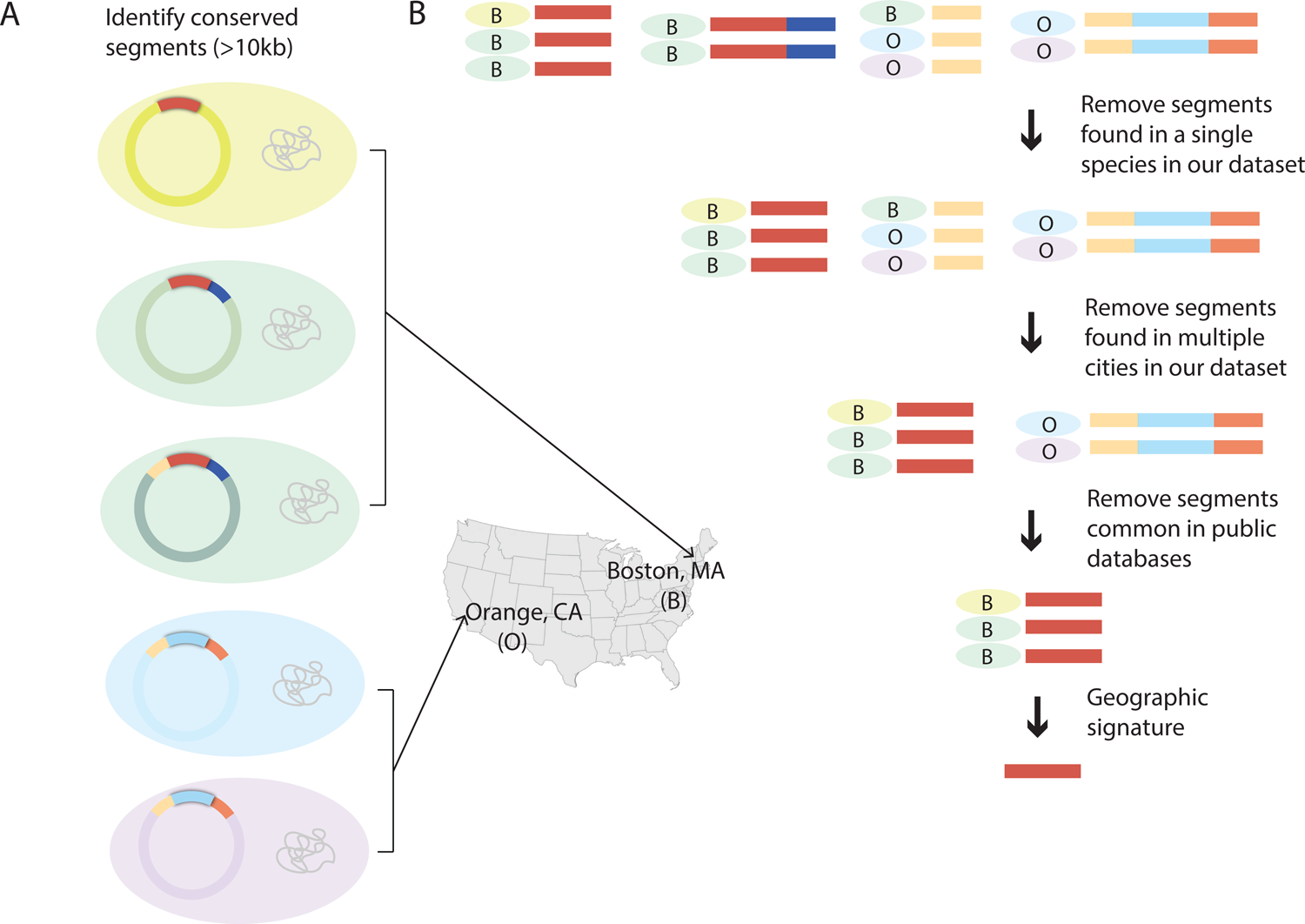
Methodology to identify highly conserved and contiguous plasmid-borne geographic signatures. **a**, Five bacterial isolates (colored by species) with conserved plasmid segments highlighted in different colors. The geographic location (city of isolation) is indicated for each. **b,** Depiction of algorithm to identify geographic signatures. Conserved segments are colored. Species and hospital of isolation are indicated for each segment.

We focused our analysis on segments that were likely horizontally transferred and present in more than one species, as interspecies transfers would be least suspected as nosocomially linked. To filter out segments likely to have been repeatedly imported into the hospital from elsewhere, we removed those present in isolates from both states represented in our study, as well as those appearing in publicly available genomes represented in the NCBI’s Nucleotide Collection database. We elected to group the Boston-based hospitals together to account for known fluidity between some hospital staff and patient populations who could interact with and share organisms from a common reservoir. Although our assemblies were highly contiguous, we took further steps to ensure that these segments represented only high-confidence regions of our assemblies and were not missed by improper assembly or scaffold breaks (Materials and Methods). Finally, we stringently screened these for uniqueness by searching against the ENA/SRA database using BIGSI [65].

This analytical framework revealed 44 *geographic signatures*, which we define as plasmid segments found in two or more species and associated exclusively with either Boston, MA or Orange, CA (Figure S9; Table S13; Supplementary Data File). These signatures represented only 2% of all segments found in multiple species, and more than half (52%) were specific to a single hospital. We found that signature prevalence was higher among CRE (23%) than CSE (6%) in our sample set, likely owing to our sampling strategy that included all CRE from hospital microbiology labs, but only a small fraction of CSE. As observed for the full set of conserved segments, many signatures exhibited substantial overlap with one another. Ten were fully nested within larger signatures, representing different conservation profiles, with each shorter signature found in more isolates than the larger signature. Signatures had a mean length of 31 kb (range 10 kb to 311 kb), and were observed, on average, 5 times (range 2 to 19), across 2 plasmid groups (range 1 to 4; 28 total), 2 species (range 2 to 4; 15 total), and 3 sequence types (range 2 to 6; 39 total).

### Geographic signatures carry important cargo for hospital survival and dissemination

We hypothesized that the 44 geographic signatures would encode functions that enable their movement and persistence within patients (i.e. colonization) and the built environment, similar to other widely conserved plasmid sequences from nosocomial bacteria. Of the 1,494 individual genes predicted within signatures, we could assign some putative function to nearly two-thirds (Figure S10-S11; Tables S14-S15). All but one of the signatures were predicted to encode functions for maintenance or dissemination of DNA into new genomic contexts or hosts, including IS elements [66], integrases or other recombinases [67], conjugation machinery [68], and plasmid uptake and maintenance apparatuses [69, 70]. The prevalence of conjugation and plasmid uptake genes expectedly pointed to the carriage of signatures on conjugative plasmids; however, one signature (Sig20) also overlapped with a predicted prophage that was situated on a circularized scaffold containing a plasmid replicon but no conjugative relaxase. Also, given their demonstrated ability to cross species boundaries, it was unsurprising that half of the signatures featured genes which were also found at high nucleotide identity (≥99%) in bacteria outside the order of *Enterobacterales* (Table S14).

As we also hypothesized, the majority (66%) of signatures featured genes predicted to encode for survival strategies against antimicrobials, including quaternary ammonium compounds used in standard disinfectants in healthcare settings [71](Figure S10-S11). Eight signatures encoded enzymatic antibiotic resistance, including examples with *bla*_KPC_, likely reflecting both our sampling strategy focused on CRE and that antibiotic resistance genes are often co-located on plasmids [72–74]. Genes coding for metal resistance were also prevalent, occurring in more than a third of signatures, and often co-occurring with genes for antibiotic resistance, including in half of signatures containing *bla*_KPC_, highlighting recent findings that metal resistance and antibiotic resistance are frequently co-selected [75, 76].

### Additional instances of localized carbapenemase spread were identified through tracing of geographic signatures

Six of the 44 geographic signatures encoded *bla*_KPC_. These *bla*_KPC_ signatures ranged in size from 18kb - 147kb (the smallest of these signatures, Sig 5.1-CP, was nested inside the largest signature, Sig 5.6-CP), and were found in 21% of all *bla*_KPC_-carrying CRE in our collection (Figure 4; Figure S9). As expected from the filters we applied to identify geographic signatures, *bla*_KPC_ signatures were distributed across multiple species (2 to 4) and sequence types (2 to 7), associated with multiple plasmid groups (1 to 3) and sometimes the chromosome, and observed across variable time spans of at least 4 years and up to 10 years (Figure 4; Table S16). Furthermore, some isolates carried more than one signature (*e.g.*, Sig1-CP and Sig4-CP), sometimes on the same plasmid.

**Figure 4.**
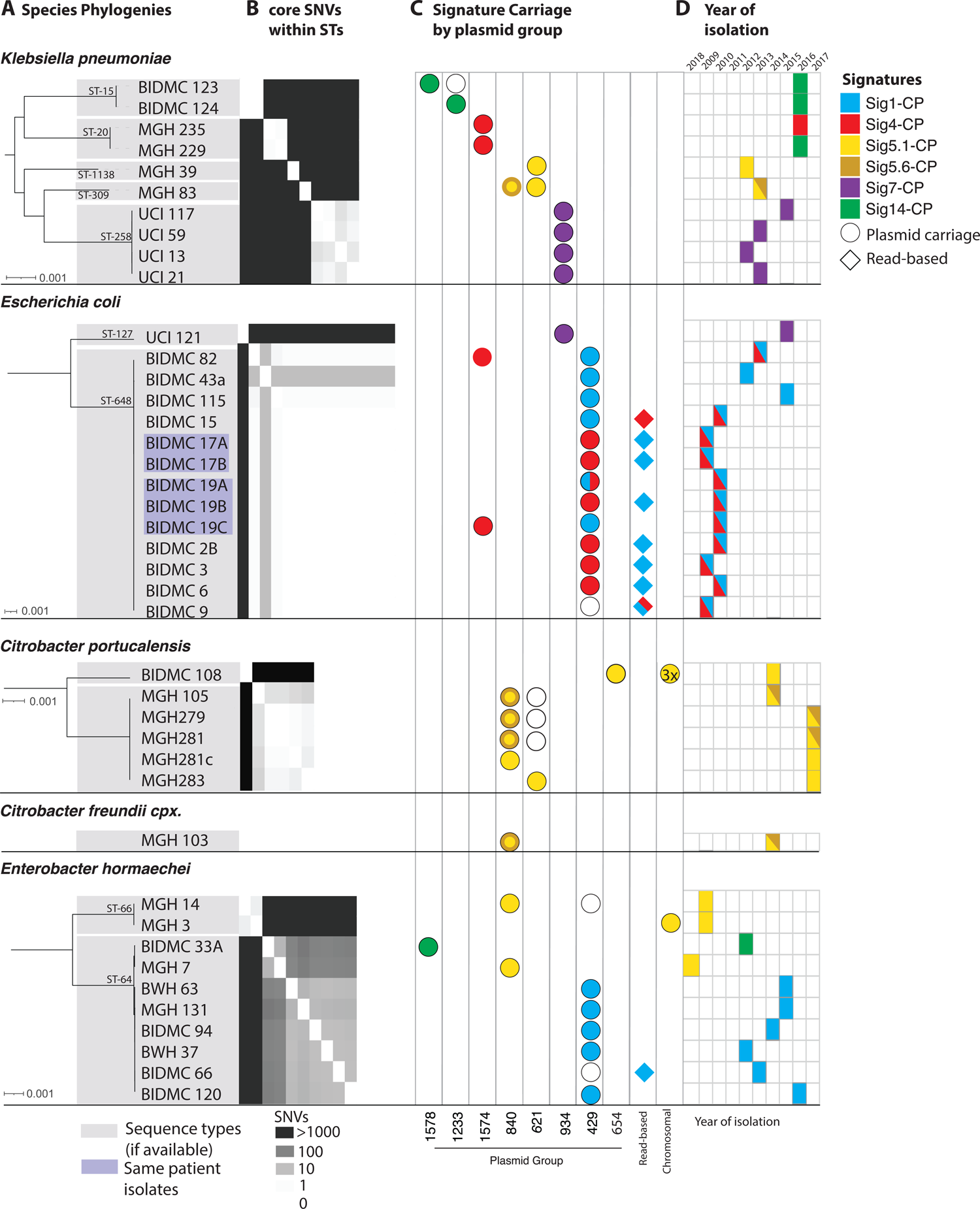
Carbapenemase-carrying signatures are found in diverse species, lineages, and plasmid backgrounds. **a,** Species phylogenies for all geographic signature-containing isolates from each species, showing sequence type (STs). **b,** Within each ST, core SNV distances are shown (heatmap). **c,** For each isolate, columns indicate plasmid content, and colored icons indicate signatures present on these plasmids. For the nested signatures 5.1-CP and 5.6-CP, a solid yellow circle indicates the presence of 5.1-CP only, whereas the yellow circle with a darker ring indicates the presence of both 5.1-CP and 5.6-CP. **d**, Year of isolation is marked for each, colored by signature content as in **c**.

Though our inclusion criteria would have allowed for as many as 100 SNVs in a 10 kb signature, we observed much greater identity, with most *bla*_KPC_ signatures varying by only two or fewer SNV differences between instances (Figure S12). This striking degree of similarity, extending well beyond the edges of the Tn*4401* elements, likely indicates a recent common ancestral source for these signatures, as well as their persistence and movement across species boundaries within a local environment. These instances of suspected local spread of carbapenemases between distantly related isolates would not have been picked up by traditional epidemiology or even standard whole genome sequence analysis in most cases.

Furthermore, many *bla*_KPC_ signature instances were harbored by isolates that were very closely related (0-10 core SNVs), indicating the ability of these signatures to spread within the hospital along with their bacterial hosts (Figure 4), in addition to their ability to move between bacterial sequence types and species.

### A multi-genus *bla*KPC-containing geographic signature is highly conserved despite rapid rearrangements of its plasmid backgrounds

Although our analysis was based on highly contiguous assemblies that combined data from short- and long-insert Illumina libraries, we sought to improve the assemblies further in order to follow the details of signature evolution and spread across plasmids and taxonomic boundaries. To do this, we re-sequenced all isolates carrying Sig5.1-CP using Oxford Nanopore Technology to generate long read sequences for hybrid assembly, followed by manual curation. Sig5.1-CP, a *bla*_KPC-3_-carrying signature, was present in the largest number of different genera and species in our prospective collection and was also found in historical isolates dating back to 2008 (Figure 5; Suppl. Table S17). In addition, Sig5.1-CP was almost exclusively found in a single Boston hospital, including within four isolates collected over a five-week period in 2017.

**Figure 5:**
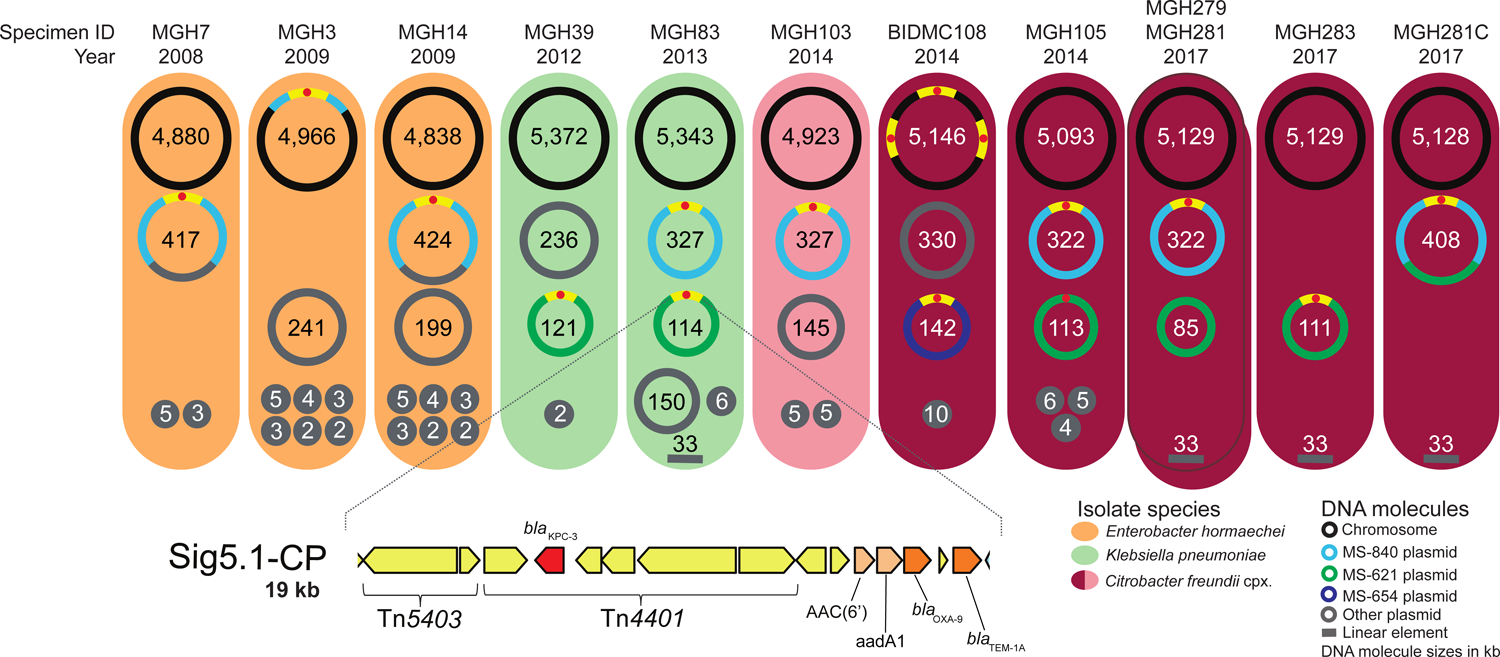
Hybrid assemblies with Oxford Nanopore long-read and Illumina short-read sequencing data for isolates harboring Sig5.1-CP. Each colored oval represents an isolate harboring signature Sig5.1-CP. Specimen IDs and the years of isolation are indicated above each oval. The DNA molecules harbored by each of the isolates are represented by circles (or lines, since a linear DNA molecule was found in four isolates) with molecule sizes indicated in kb. The location of Sig5.1-CP is shown with yellow segments; a hallmark of this signature is the truncation of transposon Tn*4401* by insertion sequence Tn*5403*. A schematic of the full 19 kb Sig5.1-CP is shown at the bottom of the figure. In this schematic, gene color corresponds to functional categorization: mobile genetic element (MGE) [yellow], carbapenem resistance [red], beta-lactam resistance [dark brown], and aminoglycoside resistance [light brown].

These four isolates were initially sequenced because they were suspected to be part of a short-lived *C. freundii* complex (later revealed to be *Citrobacter portucalensis*) clonal case cluster that differed from one another by less than four SNVs. We found that this 2017 cluster of isolates all differed by the same 12 SNVs from a 2014 isolate carrying Sig5.1-CP from the same hospital that was already part of our collection.

The complete assembly of each individual replicon unambiguously revealed the relocation of Sig5.1-CP into the chromosome and its association with three different plasmid groups (Figure 5). One of these plasmids encoded both IncP and IncH replicons, which are known for their broad host range [77, 78], likely contributing to this signature’s ability to transfer to diverse hosts and persist. Although our draft assemblies were adequate to identify the signature using our methodology, manual inspection of Sig5.1-CP boundaries in the completed assemblies revealed that this signature could be expanded to include approximately an additional 1 kb of sequence encoding a *bla_T_*_EM-1A_ beta lactamase. Manual inspection also revealed an even longer, 26kb conserved region in the 9 isolates occurring since 2012 (Figure S13). This extended signature included additional cargo that could be involved in adaptation to the healthcare environment, including genes predicted to encode mercury resistance, a Na+/H+ antiporter (possibly influencing cell viability at high pH [79]), and a multidrug efflux pump.

Despite Sig5.1-CP’s distinct genomic contexts, it was highly conserved, maintaining perfect base-level identity, with the exception of one isolate (*K. pneumoniae* MGH39 from 2012) that harbored a variant of the signature with two non-synonymous SNVs, both within the same predicted transposon.

Though possessing nearly identical chromosomes (≤3 SNVs), isolates from the *C. portucalensis* clonal case cluster had a striking variety of plasmid and chromosomal arrangements (Figure S14). While two chromosomally identical 2017 isolates also had identical plasmid profiles and Sig5.1-CP location, one of the other 2017 isolates carried the signature on a different plasmid, while the remaining 2017 isolate carried it on a co-integrate of both plasmids. Many of the rearrangements were likely mediated by IS*26* sequences (Figure S14), previously shown to drive the reorganization of plasmids through replicative transposition [80]. Although we could not precisely trace the divergence of the cluster of 2017 isolates from the 2014 isolate, multiple plasmid gains, losses, and rearrangements in the 2017 isolates appear to have occurred on the same timescale as the accumulation of SNVs.

## Discussion

Using a systematic, prospective isolate collection strategy, along with a sequencing approach that enabled highly contiguous assemblies, we found diverse mechanisms for the spread of carbapenem resistance and examples of clonal spread of both carbapenemase-carrying and porin-based CRE. We also found geographic signatures that allowed us to trace the local spread of carbapenemases across genetic backgrounds, even in cases where standard genomic epidemiology might not identify a link.

Our collection was agnostic to species and carbapenem-resistance mechanism, giving us a more complete view of patient CRE organismal and resistance mechanism diversity. As we and others observed previously [29, 30], the majority of carbapenem resistance was mediated by the presence of a carbapenemase, with the most prevalent being *bla*_KPC_, consistent with their known prevalence in the U.S. [22]. We saw that another approximately quarter of resistance could be associated with ESBL or AmpC beta-lactamases, together with at least one disrupted major porin - a resistance genotype that can be difficult to detect using targeted molecular or even whole genome sequencing analytical approaches.

Though there have been many examples of clonal spread of CP-CRE, few studies have pointed out the contribution of porin-based CRE in nosocomial CRE spread [57, 81]. Although strains with porin mutations are thought to be less successful in spreading to other patients, we observed a large clonal cluster of *K. pneumoniae* isolates with inactivating mutations in both major porins appearing in patients across a four-year period. Interestingly, the mutation in *ompK35* was the same across the isolates, whereas the mutations in *ompK36* varied. This suggested that i) the transmitted form had a single porin inactivation and, thus, may have been less impaired for long-term persistence and spread; ii) selective pressure for inactivation of both porins was able to override any fitness defect; and/or iii) the potential for yet to be identified compensatory mutations which could attenuate the fitness defect associated with mutations of these porins in this strain [82, 83]. The presence of porin-based CRE clonal clusters indicate that fitness phenotypes in such strains should be examined more closely, and that porin-based mutants should not be overlooked as contributors to clonal case clusters.

Plasmids are key to understanding the ongoing carbapenem resistance epidemic but pose challenges for tracing resistance evolution and local spread because of their diversity, widespread nature, and tendency to rearrange [31,52,54,61,84,85]. These processes are accelerated by the frequent co-occurrence of carbapenemase plasmids with other diverse plasmids within a single isolate [42]; in our dataset, three-quarters of all plasmid groups were observed to co-occur with a carbapenemase plasmid group. Furthermore, as observed here and by others [40, 42], carbapenemases are often carried by plasmids that can conjugate across species and genera [40, 42]. Our results showcase an extensive network of possible exchange points that carbapenemases (and the signatures that contain them) can use to transfer to new plasmid backgrounds and onward to new hosts. We also observed striking plasticity of plasmids. For instance, plasmid groups were rarely composed purely of carbapenemase-containing plasmids and were also widespread and not strongly associated with specific geographic locales. This flexibility is exemplified in the *C. portucalensis* clonal case cluster, in which plasmid rearrangements and carbapenemase transfer between different plasmid backgrounds was repeatedly documented for isolates that were likely to have diverged very recently and isolated only a few weeks apart.

While approaches for systematically tracing carbapenemases across species include analyzing transposons and their immediate flanking sequences [46], these regions in our collection were common and widespread, preventing us from confidently tracing carbapenemase movement. Additionally, the recently published alignment-based, pairwise screen of Evans *et al.* [64] shows promise for tracing carbapenemases; however, their clustering approach applied in this case would likely group together different Tn*4401* isoforms, and does not assess whether segments are geographically associated.

In order to overcome the limitations of these approaches and to achieve a higher level of resolution for tracing the localized spread of carbapenemases as well as other hospital adaptive traits, we developed a novel, broadly applicable, and gene-agnostic framework to identify highly conserved plasmid segments found in multiple species or lineages with strong geographic associations. Though strict filters for identifying signatures were used in the work presented here, our approach can be tuned with regard to these filters, including the ability to incorporate different levels of geographic specificity (i.e. hospital, city, etc.), phylogenetic specificity (i.e. lineage, species, etc.), and different signature lengths, up to the size of an entire plasmid.

Loosening these requirements - along with use of even more contiguous assemblies - could yield a more comprehensive profile of HGT signals, including an understanding of the possible differences in HGT occurring within versus between different species. Although we only searched for inter-species signatures in this study, we also identified intra-species instances in 8 of the 44 signatures.

While long-read technologies are typically needed to achieve the high level of contiguity necessary to examine plasmids in detail, our strategy, involving both long and short insert Illumina libraries, together with our novel approach for identifying signatures, was able to successfully identify many highly conserved signatures involved in the predicted local movement of carbapenemases across species boundaries. Although having high quality assemblies was key to identifying signatures, our method also includes searches against raw sequence reads to identify additional instances of motifs, which would make it possible to include read sets from lower quality assemblies in analysis. Improved genomic assemblies, including more long read sequencing, would likely assist in identifying and tracing signatures more completely. Furthermore, given our method’s dependence upon public databases, growth in the number of deposited sequences will provide additional resolution to discern true geographic signatures from segments that are more broadly geographically represented.

The links between unrelated isolates uncovered by geographic signatures adds to the growing recognition that there are likely to be reservoirs of *Enterobacterales* within healthcare networks that are involved in reshuffling plasmids and their signatures across organisms, and aiding in their long-term local persistence. Remarkably, over 20% of CP-CRE isolates carried at least one *bla*_KPC_ geographic signature, suggesting that a substantial fraction of patient CRE may have originated from HGT within the hospital (or hospital-proximal environments, including ambulatory and nursing facilities). Although we expected that selective pressure for maintaining carbapenem resistance would be mainly present within patients treated with carbapenems, our *bla*_KPC_-containing signatures were long-lived, each being observed in patient isolates spanning periods of four to ten years. This is likely because these signatures also carried other genes key to adaptation in hospital reservoirs. Genes co-localized with *bla*_KPC_ within our signatures that could provide functions related to persistence in the hospital environment included: i) additional antibiotic resistance genes, which could lead to their joint long-term conservation through co-selection; ii) genes conferring resistance to hospital disinfectants and metals, increasingly used in hospital touch surfaces [75, 76] and iii) conjugation and plasmid uptake machinery, both of which can amplify *bla*_KPC_ and assist its persistence and spread across isolates and species [86, 87]. In addition, the association of these signatures with plasmids from different incompatibility groups, particularly those adapted to different conditions such as temperature [88–90], appear adaptive for reservoir switching, *e.g.* from the human body to the hospital environment and back. Our views of the taxonomic distributions of signature genes suggest that some of these hospital-adaptive traits may have been recently acquired from distantly related organisms in shared reservoirs.

Our study had several limitations. Sampling of suspected hospital reservoirs, including environmental samples and those from asymptomatic carriers, would likely allow us to create a more complete view of signatures and their exchange across organisms and demonstrate the role of these reservoirs in seeding infections. Other studies, including our previous work [30], point to high levels of asymptomatic colonization, including a study that suggests that clinical testing detects only one out of nine carriers [91]. We also lacked epidemiological data about patients, which further limited our ability to quantify the extent of clonal spread. Despite the lack of epidemiological data and non-patient samples, our analysis suggested that there is persistence of resistance genes within hospital networks for years.

One additional limitation of our approach is our inability to distinguish convergent evolution or re-introductions from the community from local spread. However, due to the high degree of similarity over long stretches (10-310kb), convergent evolution for all of these signatures is unlikely. In the case of two signatures (Sig1-CP and Sig4-CP), instances that were otherwise nearly identical differed in which *bla*_KPC_ allele or Tn*4401* isoform they carried. Possible explanations other than *de novo* formation of a similar signature include mutation within a signature, or recombination taking place within the signature. The propensity for some signatures, like Sig5.1-CP, to persist for so long within a single hospital, yet to not be found more generally, including among patients treated at other nearby hospitals, strongly hints at a local reservoir, rather than re-introduction.

In conclusion, using a long-term systematic collection of isolates - including both CRE and CSE - together with high quality sequencing and a novel analysis methodology, we achieved high resolution views of mechanisms accounting for carbapenem resistance and a greater understanding of their spread across four U.S. hospitals. In addition to examples of clonal spread of CP-CRE and porin-based CRE, we observed the sharing of plasmid segments containing hospital adaptive traits - including carbapenemases - circulating among local diverse bacterial populations over long timeframes. Within these segments and the plasmids harboring them, we observed intermolecular rearrangements over short timeframes, underscoring the complexity entailed in tracing the movement of plasmids and their component parts. However, our long-term surveillance strategy, high quality assemblies, and novel methodology for identifying geographic signatures revealed previously unsuspected links between CP-CRE (as well as other organisms across these hospitals) that helped to clarify the epidemiology of antibiotic resistance spread and persistence in these healthcare networks.

## Materials and Methods

### Isolate collection and drug susceptibility testing

Our sample collection represented a continuation of our previous prospective study, conducted in 2012-2013 [30]. Between December 2013 and November 2016, we collected additional samples from symptomatic patients at Beth Israel Deaconess Medical Center (BIDMC), Brigham and Women’s Hospital (BWH), and Massachusetts General Hospital (MGH) in Boston, MA, and the University of California Irvine (UCI) Medical Center in Orange, CA. We collected all isolates from clinical samples sent to these hospitals’ clinical microbiology laboratories that, using the laboratories’ standard operating procedures [30], were identified as *Enterobacterales* with a meropenem minimum inhibitory concentration (MIC) ≥ 2 µg/mL, the threshold that we used to define resistance throughout our analysis. We thus included samples categorized as both *intermediate* (2 µg/mL) and *resistant* (≥ 4 µg/mL) by the Clinical and Laboratory Standards Institute interpretative criteria [92]. For each resistant isolate, we also collected date- and species-matched meropenem susceptible isolates. UCI submitted two carbapenem susceptible isolates per resistant isolate; the other three hospitals submitted a single susceptible isolate per resistant isolate. In total, we collected 347 new isolates (146 resistant and 201 susceptible isolates; one isolate per patient). Isolates whose meropenem resistance phenotype was discordant with the genotype were retested using a previously validated automated digital dispensing method [93] adapted from standard broth microdilution procedures [92]. Meropenem was tested against bacterial isolates at doubling dilution concentrations from 0.016 to 32 µg/mL. All isolate cultures are available from the corresponding author upon reasonable request.

Combining our newly collected samples from December 2013 - November 2016 with the 261 *Enterobacterales* isolates that we previously sequenced in 2012-2013 from these four hospitals [30] resulted in a total of 608 isolates (of which 274 were meropenem-resistant). The 261 previously reported isolates consisted of the retrospective *Boston Historical Collection*, consisting of 49 isolates collected between September 2007 and July 2012, including 17 isolates grouped into six sets of same-patient isolates, and 74 CRE and 138 carbapenem susceptible *Enterobacterales* (CSE) isolates collected prospectively (including 11 isolates from 3 sets of same-patient isolates). In total, our prospective collection included 559 isolates collected between 2012 and 2016.

Institutional review board (IRB) approval was granted by the Massachusetts Institute of Technology Committee on the Use of Humans as Experimental Subjects. Samples were collected under study approvals of the IRB committees of the participating institutions: Mass General Brigham (covering both MGH and BWH); Beth Israel Deaconess Medical Center, Boston; and University of California, Irvine.

### Genome sequencing, assembly, and annotation

*<u>Illumina sequencing and assembly</u>*. We prepared whole-genome paired-end and mate-pair libraries for 347 isolates and paired-end sequenced them as previously described [30] on Illumina HiSeq 2500 or HiSeq X sequencers. Genome assembly and annotation were carried out as previously described, using the Broad Institute’s prokaryotic annotation pipeline [30]. Sequencing reads and assemblies were submitted to GenBank under bioproject PRJNA271899.

*<u>Long-read sequencing and assembly</u>*. We selected a subset of 19 isolates for further sequencing using Oxford Nanopore Technology (ONT). For the 12 isolates containing signatures Sig5.1-CP or Sig5.6-CP, we used the Oxford Nanopore Rapid Barcoding kit (SQK-RBK004) and ran the samples on a Minion (Oxford Nanopore Technologies Ltd, Science Park,UK). For an additional 7 isolates, 600 nanograms of DNA from each sample were used as input into the Oxford Nanopore 1D ligation library construction protocol (SQK-LSK109) following the manufacturer’s recommendation. Samples were barcoded using the Native Barcoding Expansion 1-12 kit to run in batches of between 1-4 samples per flow cell on a GridIon (Oxford Nanopore Technologies Ltd, Science Park, UK).

Oxford Nanopore (ONT) reads were demultiplexed using Deepbinner (v0.2.0)[94], trimmed of any remaining adapter using Porechop (v0.2.3), and subsampled to approximately 50x depth of genome coverage. Illumina reads were trimmed of adapter using Trim Galore (v0.5.0) and subsampled to approximately 100x depth of genome coverage. Two Unicycler (v0.4.3 or v0.4.4, with default settings) [95] hybrid assemblies were generated for each sample, one assembly combining the Illumina 100x data set with the 50x subsampled ONT data set and another assembly combining the Illumina 100x data set with the full set of ONT reads (if > 50x).

ONT reads were aligned to Unicycler contigs using minimap2 (v2.15) [96]. Illumina reads were aligned to Unicycler contigs using bwa mem (v0.7.17) [97], and the resulting alignments were input to Pilon (v1.23) [98] for assembly polishing. Contigs were screened for adapter sequence and then input to GAEMR (https://github.com/broadinstitute/GAEMR) which produced chart and metric tables for use in manual assembly analysis process. Hybrid assemblies were annotated as above. Reads were submitted to SRA under bioproject PRJNA271899.

*<u>Annotation of resistance genes.</u>* We annotated our assemblies for the presence of resistance genes as done previously [30], but we queried predicted genes with an updated set of antimicrobial resistance databases using BLASTn in megablast mode [99]: i) our database of carbapenem-hydrolyzing beta-lactamases [30]; ii) ResFinder [100], downloaded January 22, 2018; and iii) the antimicrobial resistance database of the National Database of Antibiotic

Resistant Organisms (https://www.ncbi.nlm.nih.gov/pathogens/antimicrobial-resistance/, downloaded January 22, 2018). Genes with hits to any of these databases (E-value < 10^-10^; query coverage ≥ 80%) were annotated with the name of the antibiotic resistance gene with the highest BLAST bit score; matches to multiple databases were resolved with the same order of precedence with which the databases are listed above (Tables S7 and S18). A subset of beta lactamases were designated as ESBLs by our pipeline for identifying resistance genes (Table S5), together with additional evidence from the literature [101].

In order to identify defects in porin sequences contributing to carbapenem resistance, we first identified all porin genes by sequence similarity search [99]. We queried our set of predicted genes against a database of *ompC* and *ompF* reference sequences [30], retaining the best hit with E-value < 10^-10^ and ≥ 80% coverage of the reference sequence. We also searched for matches in the full sequences of the assemblies, in order to identify genes or gene fragments that were not part of the predicted gene set. Within the resulting set of predicted porin genes, we then identified mutations by comparing them to the best matching reference sequence with MUMmer [102]. Additionally, we pinpointed porins disrupted by insertion sequences by submitting porin sequences and their promoter regions (500 bp upstream) to the ISfinder [103] BLAST facility, retaining hits with E-value < 10^-10^. We regarded a porin gene as disrupted if i) no BLAST hit for the gene was produced; ii) the best matching predicted gene contained < 90% of the reference sequence; iii) a frameshift mutation affected 30 codons of the gene or more; or iv) an insertion sequence was found disrupting the porin gene or up to 300 bp upstream.

In order to search for evidence of genotypic resistance that may not have been captured in our assemblies, we applied ARIBA [104], a read-based gene search tool leveraging local targeted assembly, with a database comprising carbapenemases and common ESBL genes. We considered ARIBA calls with one non-synonymous mutation or less, and a coverage of 100% (Table S19).

*<u>Plasmid Annotations</u>*. We used MOB-suite (database version from January 2, 2019) [105] to identify plasmid scaffolds, plasmid replicon types, and relaxase types. MOB-suite is a bioinformatic tool that predicts and categorizes plasmid scaffolds into discrete “plasmid groups” by clustering input scaffolds with reference plasmids if their estimated ANI [106] is at least 95%. To supplement MOB-suite’s plasmid replicon database, we added 111 additional replicons from the PlasmidFinder database [107]. The tool either assigned a reference plasmid group to each plasmid scaffold or called it “novel” if there was no match. In each assembly, plasmid scaffolds with the same plasmid group were predicted to be part of the same plasmid and grouped (but “novel” scaffolds were not grouped). While the largest scaffold to be assigned a plasmid group was 481 kb in size, longer (500 kb to 5.6 Mb) scaffolds marked as “novel” were inspected and frequently found to be chromosomal and to contain plasmid replicon and relaxase genes, suggesting that they contained integrative plasmids or similar elements [108]. For this reason, 162 “novel” scaffolds with length > 500 kb were not assumed to be part of plasmids.

### Comparative genomics

*<u>Orthogroup clustering and construction of multi-species phylogeny</u>*. In order to identify genes shared between isolates, we performed orthogroup clustering for our entire set of 608 genomes using Synerclust [109], a tool which provided the high level of scalability needed for this large set of genomes, and also leveraged the syntenic organization of genes to help in defining orthogroups. We generated a final set of orthogroups using an iterative, two-step process. First, we ran Synerclust with an approximate, *k*-mer based input dendrogram generated by i) k-merizing our set of genomes with a k-mer size of 15; ii) computing a similarity matrix using the Jaccard index [106] to compare each pair of genomes in our dataset; and iii) computing a dendrogram with the neighbor joining tree algorithm [110] contained in the ape [111] package (v5.0) of R [112] (v3.4.0). We then ran SynerClust with default parameters in order to identify ortholog clusters. We produced a codon-based multiple sequence alignment for each single-copy core gene using Muscle [113] and produced a concatenated alignment of all genes by extracting alignment columns without gaps. We then computed a phylogenetic tree using Fasttree [114] with default parameters, which we then used as input for a second iteration of Synerclust. The orthogroup output from the second iteration of Synerclust was used to establish the final single copy core gene set to be used for downstream analysis, including construction of a single copy core alignment (as in the first iteration). This final alignment spanned 676,371 nucleotide sites, of which 348,152 were variable, and was used to generate a phylogenetic tree using RAxML [115] (v7.3.3), using rapid analysis of 1,000 bootstrap replicates. To generate phylogenetic trees containing only subsets of isolates, we used PareTree (http://emmahodcroft.com/PareTree.html).

*<u>Species identification</u>*. We used average nucleotide identity (ANI) to obtain species designations for each isolate. For each pair of isolates, we used alignments of all shared genes (using orthogroup clusters) to compute ANI [116]. We compared to reference assemblies obtained from the NCBI taxonomy browser (https://www.ncbi.nlm.nih.gov/taxonomy) to obtain species designations.

*<u>Construction of species-specific multiple sequence alignments of the core genome.</u>* In order to construct more detailed, species-specific SNV-based phylogenies, for each species we selected the assembly with the smallest number of contigs as a reference. Then, we produced alignments for both the short paired-end and long-insert mate pair sequencing reads of each isolate using bwa mem [97] and sorted the alignments with Picard SortSam (v2.20.6; http://broadinstitute.github.io/picard). Finally, we used Pilon (v1.23) [98] in order to call variants.

We produced multiple sequence alignments based on the variant calls for all isolates of each species, excluding alignment positions with insertions or deletions. The number of nucleotide sites in these alignments ranged from 2,931,929 sites for *E. coli* to 5,048,275 for *K. oxytoca* (Table S20).

*<u>Computing phylogenetic trees after removing effects of recombination</u>*. To construct more accurate species-specific phylogenies using our SNV-based alignments, we used ClonalFrameML (v1.11) [117] to identify and remove alignment regions with evidence for recombination. We ran ClonalFrameML with default parameters and 100 bootstrap replicates, using an input phylogenetic tree generated with FastTree (v2.1.3) [114], and 100 bootstrap replicates. We produced a recombination-removed multiple sequence alignment by removing any site from the species-specific alignment in which recombination was detected in at least one isolate. The number of nucleotide sites in the resulting alignments ranged from 593,538 for *E. coli* to 4,978,737 for *K. oxytoca* (Table S20). Using these alignments, we used RAxML [115] (v7.3.3) with 1,000 rapid bootstrap replicates to generate final phylogenies. We also used these alignments to calculate core SNV distances between each pair of isolates.

*<u>Determination of Lineages and Sequence Types</u>*. We computationally determined lineages in each species using the recombination-removed phylogenetic trees. We assigned isolates to the same lineage if they were connected by a path consisting entirely of branches with a length of 10^-4^ substitutions per nucleotide site or less. Sequence types were computationally determined as before [30]. In brief, sequence types were determined using our Broad pipeline for determining sequence types. In brief, this script uses BLAST to compare the assembly against a database of sequences from pubMLST [118] using a 95% threshold in order to predict the sequence type. For the isolates belonging to each sequence type, we identified the lineage most commonly assigned to the members of this sequence type; this mapping produced the correct lineage in 87% of isolates. Conversely, for the isolates belonging to each lineage, we identified the sequence type most commonly assigned to members of this lineage; this mapping was correct for 98% of isolates. Lineages corresponding to the following sequence types were considered *high-risk* [43]: *E. coli* ST38, ST69, ST131, ST155, ST393, ST405, and ST648 and *K. pneumoniae* ST14, ST37, ST147, and ST258.

*<u>Assessment of established genetic markers to trace local movement of bla_KPC_.</u>* We characterized Tn*4401* structural variants and isoforms, as well as target site duplication (TSD) flanking sequences of Tn*4401* using TETyper [46] for the 608 samples included in our study. We analyzed 5 bp surrounding the three most common Tn*4401* isoforms (Tn*4401*a, Tn*4401*b, and Tn*4401*d) in our assemblies.

### Identification of geographic signatures

We developed the ConSequences software suite to identify nearly identical ≥ 10 kb segments conserved between two or more plasmids (Figure S7-S8). Along with the open-source code, a test dataset for running the three primary programs in ConSequences can be found on its GitHub repository (https://github.com/broadinstitute/ConSequences), consisting of the twelve hybrid assemblies constructed using Illumina and ONT sequencing for isolates found to harbor the geographic signature Sig5.1-CP.

*<u>Selection of plasmid sequences</u>.* To construct the database of plasmid sequences that we searched for geographic signatures, we included all scaffolds between 10 kb and 500 kb that were not classified as chromosomal by MOB-suite [105]. We excluded scaffolds classified as plasmidic and longer than 500 kb, since we found these to be often misclassified (*Plasmid annotations*). Circular and complete representations of plasmids [119, 120] were determined when a plasmid scaffold showed both significant overlap between its ends (e-value < 10^-5^) and had at least five mate pair sequencing reads bridging scaffold ends.

*<u>Identification of highly conserved 10 kb windows shared across pairs of plasmid backbones</u>*. All plasmid-predicted scaffolds were aligned in a pairwise manner using BLASTn in megablast mode. In order to account for circularity of complete plasmids [119, 120], we duplicated 10 kb from the beginning of the scaffold and appended it to its end. For each scaffold, a sliding window approach, with a window size of 10 kb and a step size of 100 bp, was applied to identify highly conserved and contiguous windows shared with at least one other scaffold, where matches were required to exhibit ≥ 99% identity and coverage through single high scoring pairs (HSPs). A 10 kb window size was selected for the analysis because i) the vast majority (98.7%) of HSPs with identity ≥ 99% were shorter; thus, 10 kb and longer sequences were outliers and hypothesized to share a recent ancestral origin; and ii) it allowed us to capture whole operons or large transposable elements and their surrounding contexts. For example, isoforms of *Tn4401* typically span around 10 kb, are well conserved, and are often found on different plasmid backbones [30, 46].

*<u>Delineating boundaries of shared segments between plasmids</u>.* We developed a novel algorithm to identify the boundaries of shared segments spanning multiple adjacent windows along a reference plasmid scaffold by first traversing blocks of adjacent windows in the forward direction, and then repeating the process in the reverse direction. For each 10 kb window in the series, the *focal window*, we checked whether downstream windows showed conservation in the same set of scaffolds as the focal window, tracking how far the *segment* could potentially be expanded. This procedure was then repeated in the reverse direction for the same series of windows. After potential segments were identified from both forward and reverse traversals, they were merged if they exhibited overlap in coordinates and shared conservation in a common set of scaffolds (Figure S7). As identical segments were often obtained by using different reference scaffolds, we used CD-HIT to cluster sequences with ≥ 99% global identity and ≥ 95% coverage of both sequences. Representative segments were selected from each cluster by maximizing for the number of samples segments were found in.

*<u>Filtering shared segments to identify geographic signatures</u>*. In order to identify signatures, we filtered for segments which had broad host range and exhibited geographic association. Starting with the set of segments conserved across multiple species, we identified those which were found exclusively in isolates from a single city (Boston, MA or Orange, CA). These segments were then screened for uniqueness against NCBI’s Nucleotide Collection database (nt; downloaded in December 2019), using thresholds of 98% identity and 95% query coverage. Hits matching samples in NCBI sourced from the same city were retained as potential geographic signatures.

The presence of assembly errors, including incorrect copy counts for tandem repeats, could lead to the incorrect association of segments with geographies. Thus, we checked whether any of the potential geographic signatures contained such tandem repeats using Pilon [98]. Paired-end library sequencing reads from each isolate found to harbor a signature were aligned to the signature’s reference sequence using bwa mem (v0.7.17 with default settings) [97]. Then Pilon (v1.23) [98] was run with the options: “--vcf --fix all,breaks --mindepth 5.” Instances which triggered a fix break report with the flag“TandemRepeat”, indicating the segment likely contained a tandem repeat motif, were identified and removed to ensure geographic association was not driven by faulty estimation of the tandem repeat motif’s copy count.

We next performed a more comprehensive assessment of uniqueness for each of the signatures using BIGSI [65], searching against all raw sequencing read sets in a snapshot of the ENA/SRA database taken on December 2016, contemporaneous with the most recent isolation dates for the 608 samples in this study. The SRA/ENA snapshot provided a broader database (455,632 read sets) compared to our original screening against the nt database, which included only a subset of assemblies available in the nt database and did not account for bacterial samples with sequencing data but no assembly. To perform this search, we used a sliding window (2 kb window; 1 kb step) across each signature to identify read sets containing at least 99% of all *k*-mers for each window. Matching read sets were downloaded from EBI’s ENA database and further searched using a *k*-mer based methodology (described below) to more stringently assess whether they harbored any of the geographic signature sequences.

*<u>Searching for additional instances of geographic signatures directly in raw Illumina sequencing read sets.</u>* It is possible that instances of geographic signatures were missed in our dataset of predicted plasmid segments since i) not all assemblies contained finished, circular representations of plasmids and ii) chromosomal scaffolds were not accounted for in our original search for signatures.

In order to recover missing signature instances, we searched the raw sequencing read sets against each multi-species geographic signature (Suppl Figure S8). First, we created reference guided multiple-sequence alignments for each geographic signature from all assemblies that contained that sequence. For each isolate, raw sequencing reads from both paired-end and mate-pair libraries were then downsampled to ∼100x. All 31-mers that were observed at least five times in each read set were next compared to each signature multiple-sequence alignment. A signature was considered present when all 31-mer windows along the multiple-sequence alignment had a corresponding match in the sample’s set of 31-mers. As slight variations can exist between instances of a signature in the multiple sequence alignment, a sample only needed to possess one of the possible 31-mers. Windows encompassing small deletions, insertions, or missing characters were ignored.

To further validate additional signature instances identified by the *k*-mer-based approach, we aligned a representative sequence for each signature to the draft assembly of the sample the sequence was extracted from using BLASTn. Hits that achieved identity > 98%, signature coverage > 95% were retained for downstream analysis and were often captured from chromosomal scaffolds that were not part of the plasmid fraction that was originally analyzed or were missed due to assembly fragmentation. To prevent incorporating false positives into our analysis, we excluded instances where the assembly included only part of the respective signature that was embedded fully within a scaffold or had no significant alignment to a sample’s assembly. To further refine the list of geographic signatures, we also checked whether smaller signatures nested within larger signatures were found in the same set of isolates (had identical conservation profiles). For such cases, we excluded the smaller nested signature from consideration.

*<u>Functional annotation of signature gene content.</u>* In order to characterize the diversity of functions encoded by plasmids and signatures, we clustered predicted protein sequences from all plasmid scaffolds larger than 10 kb using CD-HIT [121] with the parameters c=0.95, aS=0.9, and aL=0.9 (95% identity and 90% subject and query coverage). For each cluster, a representative protein was annotated by: i) using the Broad Institute’s prokaryotic annotation pipeline [30]; ii) transferring annotations from BLAST matches (≥ 90% identity and ≥ 80% coverage) to NCBI RefSeq’s non-redundant database of bacterial proteins (BacNR); and iii) transferring protein-domain annotations from Pfam [122]. Phages were identified using ProphET [123]. To further refine our annotations of the subset of the genes found in signatures, we used keyword searches on these combined annotations, together with other gene-family-specific tools to identify genes within six broad functional categories of interest (Tables S14 and S15).

<u>Antibiotic resistance genes</u> were predicted using methods described above (*Annotation of resistance genes*). <u>Chemical and heavy metal resistance operons</u>, providing resistance to mercury, arsenic, tellurium, nickel, and copper, were identified by keyword searches within our combined annotations. Operons were considered when they were composed of three or more functionally relevant genes located in close physical proximity to each other. <u>Genes involved in efflux or response to stressors</u>, including stressor efflux and transport (*e.g. silE*, *crcB*, *fieF*, *sugE*) and response genes (*e.g. dnaJ, usmG, frmR*) were identified by searching for keywords within our combined annotations. BLAST alignment [124] of proteins to representative transporter proteins in TCDB [125] (e-value < 10^-10^) was also used to flag additional proteins which might be involved in efflux, and such proteins were further examined through alignment to NCBI’s comprehensive NR database. <u>Conjugation machinery,</u> notoriously difficult to identify and differentiate from other type IV secretion systems [126], was flagged using MacSyFinder tool [127] with CONJscan HMMs [68, 128]. To improve sensitivity, we also classified additional genes as likely related to conjugation machinery based on keywords found in our combined annotations. <u>Plasmid uptake machinery</u> included type I and II toxin-antitoxin systems [69] and anti-restriction proteins [129, 130]. Toxin and antitoxin genes were predicted using HMMer v3 [131] with HMMs from the TAsmania database [70] (e-value <10^-5^) and filtered for the likelihood of representing a true toxin/antitoxin system through manual assessment of annotations. Anti-restriction proteins were identified by using keyword searches. <u>Genes associated with mobile genetic elements</u> included transposases, integrases or other recombinases, and homing endonucleases. Insertion elements and transposon genes were identified using ISFinder [103], as described above (*Annotation of resistance genes*). We found additional instances by searching for the keywords ‘transposases’ and ‘IS’ in general annotations together with manual inspection. Genes corresponding to integrases or alternate recombinases as well as homing endonucleases were also identified using keyword searches and manual validation.

To assess whether any genes found in signatures originated from sources outside the order of *Enterobacterales*, we aligned the nucleotide sequences of each gene to NCBI’s Nucleotide Collection database (nt; July 2020) using BLASTn [124]. For each gene, the top 100 hits in nt were selected based on bitscore and then filtered to ensure they matched the query gene at 99% identity and 90% coverage. Next, the taxonomic information of each target sequence was extracted from the Entrez database using Biopython to enable the calculation of what percentage belonged to bacteria from outside *Enterobacterales*.

## Statistical analysis

For assessments of statistical significance, a *p*-value threshold of 0.05 was used. Statistical significance of the differences in genome sizes and plasmid counts for CRE vs. CSE, as well as for the co-occurrence of plasmids groups, was assessed using a two-sided Wilcoxon rank sum test. The statistical significance of the differences in relaxase carriage in plasmids carrying carbapenemases vs. other plasmids were assessed using a two-sided Fisher’s exact test. The statistical significance for the increase in proportion of ESBLs over time was calculated using a regression slope test.

## Code availability

Computer code used for the analysis of our data can be downloaded from https://github.com/broadinstitute/ConSequences. A yaml file is provided for installation of the software and its dependencies through creation of a Conda virtual environment. ConSequences is written in Python3 and made available under the open-source license BSD3.

## Supporting information

Suppelementary Figures S2-S14

Supplementary results

## Declarations

### Ethics approval and consent to participate

Not applicable.

### Consent for publication

Not applicable.

### Availability of data and materials

The dataset analysed during the current study is available in the National Center for Biotechnology Information (NCBI) repository under BioProject PRJNA271899 (https://www.ncbi.nlm.nih.gov/bioproject/?term=prjna271899).

### Competing interests

B.J.W. is an employee of Applied Invention (Cambridge, MA). No other authors declare competing interests.

## Funding

This work was supported by NIH grant U19AI110818 to the Broad Institute, by NIH grant R33AI119114 to JEK. and by NIH training grants T32HD055148, T32AI007061, 1K08AI132716, and Boston Children’s Hospital Office of Faculty Development Faculty Career Development fellowship to TB-K, and by Applied Invention, LLC to BJW. The contents of this publication are solely the responsibility of the authors and do not necessarily represent the official views of the funders.

## Authors’ contributions

Study design. ALM, ABO, DCH, DTH, LAC, SSH, JEK, RPB, and AME

Study coordination. SBC, WH, MLD, CEB, GMG, DK, EMP, MJF, and AME

Assays performed. TB-K, PM, LLH, JQ, and JIW

Data analysis. RS, ALM, BJW, CJW, TPS, and AP

Consultation and supervision of analyses. ALM, TB-K, CJW, PM, SKY, SK, DCH, ESS, CAC, SSH, JEK, VMP, RPB, and AME

Prepared the original draft. RS, ALM, BJW, CJW, AP, and AME

Review and approval of the final manuscript was provided by all authors.

## Acknowledgements

The authors would like to gratefully acknowledge the patients from whom study isolates derived. The authors would also like to thank Thomas Abeel, Paul Cao, Gustavo Cerqueira, Christoph Ernst, Michael S. Gilmore, Zamin Iqbal, Jonathan Livny, Amy Mathers, and Noam Shoresh for helpful discussions as well as members of the Broad Bacterial Genomics group. We also thank members of the Broad Technology Labs and Microbial ‘Omics Core for their assistance with data generation.

## Supplemental Figures

Provided as two separate multi-page PDF files:

2021_CRE_GenomeMed_SupFigures_S1.pdf and

2021_CRE_GenomeMed_SupFigures_S2-S13.pdf.

**Figure S1.** Phylogenetic tree for each of the 15 species in our collection with at least five representatives. Below the phylogenies, colored strips indicate resistance mechanisms and hospital of isolation for each isolate. MGH: Massachusetts General Hospital, Boston, MA; UCI: University of California, Irvine, CA; BIDMC: Beth Israel Deaconess Medical Center, Boston, MA; BWH: Brigham and Women’s Hospital, Boston, MA.

**Figure S2. Resistance mechanisms were diverse, with many shared across species.** The numbers of resistant isolates by resistance mechanism (different colors) are depicted by species.

**Figure S3. Carbapenemase-carrying isolates tended to have a higher minimum inhibitory concentration than those with other resistance mechanisms.** Minimum inhibitory concentrations (MICs) of the isolates in our collection are depicted, stratified by mechanism of resistance (different colors).

**Figure S4. High level of diversity and phylogenetic range among predicted plasmids.** This plot displays the 215 plasmid groups (rows) contained in all 506 isolates (columns) for which plasmids were predicted. Plasmids with carbapenemases are indicated in red, and plasmids without carbapenemases are indicated in white. Isolates are ordered phylogenetically, while the plasmid groups are ordered by the number of genera in which they occurred and clustered.

**Figure S5: Plasmids of diverse groups carried carbapenemases and were found in different species, and hospitals.** The number of plasmids from groups for which we found at least four instances is shown. Groups with plasmids that carry carbapenemases (CPs) are depicted on a grey background, while those not observed to carry carbapenemases are shown on a white background. **a,** Plasmid instances colored by carbapenemase carriage. **b**, Plasmid instances colored by genus. **c**, Plasmid groups colored by hospital of origin.

**Figure S6: Limited tracing of carbapenemase localized spread using Tn*4401* isoforms and their immediate flanking sequences. a,** Number of instances for Tn*4401* isoforms; and **b,** combinations of the three most common Tn*4401* isoforms with their 5 bp flanking sequences which were found in multiple isolates from our study. Colors indicate the proportion of instances found in each of the four hospitals. The asterisk indicates forms that were found in multiple species.

**Figure S7: Method for the delineation of segments shared between plasmids.** *ConSequences* identifies the boundaries of conserved segments spanning multiple 10 kb windows which can be found across multiple (> 2) isolates through assessment of conservation profiles across adjacent windows along reference scaffolds. **a**, Each bar depicts a 10 kb window highlighted by sliding window analysis as being conserved in multiple scaffolds. These bars are ordered along the reference scaffold positionally (x-axis) and the height of bars corresponds to the number of scaffolds in our isolate assemblies that have a highly similar match (≥ 99%) to the 10 kb sequence on the reference (colored by genus). **b**, Using a custom algorithm (*Materials and Methods*), segments ≥ 10 kb were delineated along the reference scaffold based on conservation profiles across multiple adjacent windows.

**Figure S8: Workflow to identify geographic signatures.** The number of plasmid segments that were retained after sequentially applying different filters to identify 44 geographic signatures is shown (*Materials and Methods*). The number of plasmid segments carrying carbapenemases (CPs) is provided in red.

**Figure S9. Signatures were present across diverse sequence types.** In this heatmap, each row corresponds to a unique signature, and each column corresponds to a sequence type (ST). The shading represents the percentage of signature instances belonging to different taxonomic lineages, species or ST. The bar plot to the left of the heatmap depicts the number of isolates containing each signature, highlighting their prevalence across different hospitals.

**Figure S10. Signatures carried genes important for hospital adaptation and signature mobility. a,** Each row in the heatmap corresponds to one of the 44 geographic signatures. Groups of signatures that nest into each other are separated by horizontal dashed lines. The predicted functions of 1,494 genes within our 44 signatures were categorized into five major functional categories, unless they fell outside of these categories (*other*) or no gene function could be predicted (*hypothetical)*. The coloring of the heatmap indicates the percentage of genes of each signature that are assigned to a particular category. The identifiers of carbapenemase-carrying signatures are shown in red type and suffixed with *-CP*. **b.** Number of genes in each signature.

**Figure S11: Details of signature content.** Schematics are shown for the gene content of each signature, including the five with *bla*_KPC_. Genes are colored according to broad functional categorizations (*Materials and Methods*).

**Figure S12: Signatures were highly conserved and likely derived from a common ancestral sequence.** The number of confident, unambiguous single nucleotide variants (SNVs) differentiating signature instances was calculated for each of the 44 geographic signatures, through comparison of each instance to the signature’s representative sequence using Pilon [98]. **a**, Number of isolates carrying each signature. **b**, Box plot of SNV frequencies. SNV frequencies were calculated by normalizing the count of SNVs between each signature instance and the reference sequence (**c**) with the signature’s length.

**Figure 13. Geographic signatures with *bla*_KPC_ can occur in multiple configurations across several species and plasmid groups.** The heatmap on the left indicates the presence of signatures Sig5.6-CP and Sig5.1-CP, and the alternate boundaries of the latter, across the twelve isolates found to harbor the signature(s). The gene content of each signature is shown on the right.

**Figure S14. Geographic signatures with *bla*_KPC_ occurred in multiple configurations across several species and plasmid groups. a,** The core genome single-nucleotide variants (SNVs) and plasmid and geographic signature carriage of five nearly identical *Citrobacter portucalensis* isolates is shown. **b,** Alignment of the MS-621 plasmids carried by all isolates **a**. Two of these plasmids carry Sig5.1-CP, indicated with the bright yellow bars and triangles. The locations of the *bla*_KPC_ and of insertion sequence IS26 are indicated with red and grey rectangles, respectively. Inversions in the alignment are indicated with orange connector lines; matching regions are indicated with green connector lines. In one isolate, plasmids MS-840 and MS-621 cointegrated, which is indicated by blue alignment flanks.

## Supplementary Tables

Provided as a single Excel spreadsheet, where each supplementary table corresponds to a different tab: 2021_CRE_GenomeMed_Supplementary_Tables.xlsx

**Supplementary Table S1**: Isolates in our dataset

**Supplementary Table S2**: Accessions for Illumina-ONT hybrid assemblies

**Supplementary Table S3**. Differences in the core genome between same-species isolates from the same patient

**Supplementary Table S4**. Genome assembly statistics

**Supplementary Table S5**: List of genes identified as ESBL or AmpC

**Supplementary Table S6**. Disruptions in ompC/OmpK36 porin gene

**Supplementary Table S7**. Disruptions in ompF/OmpK35 porin gene

**Supplementary Table S8**: Carbapenemase carrying isolates.

**Supplementary Table S9**: Pairs of closely-related CSE and CRE isolates, where the CRE carried a carbapenemase not found in the CSE

**Supplementary Table S10**: Cluster of related Klebsiella pneumoniae isolates with double-porin mutations

**Supplementary Table S11**. Identification of circular plasmids (>2 kb)

**Supplementary Table S12**. MOB-suite plasmid group predictions

**Supplementary Table S13**. 44 multi-species geographic signatures specific to a single city.

**Supplementary Table S14**. Functional annotation of genes present in 44 signatures.

**Supplementary Table S15**. Prevalence of functional categories across the 44 signatures.

**Supplementary Table S16**: Isolates which carry a bla_KPC_ -containing signature.

**Supplementary Table S17**: Genomic background of instances of Sig5.1-CP and Sig5.6-CP

**Supplementary Table S18**: Genomic location of the resistance genes

**Supplementary Table S19**: Read-based identification of carbapenemases in resistant isolates for which no resistance mechanism was found in the assemblies

**Supplementary Table S20**: Size of species-specific core-genome alignments

## Additional Supplementary Documents

Supplementary Results: Supplementary Results.

Supplementary Data File: Representative sequences of 44 geographic signatures.

