## Supplementary material for "Inter-species geographic signatures for tracing horizontal gene transfer and long-term persistence of carbapenem resistance": Suppelementary Figures S2-S14

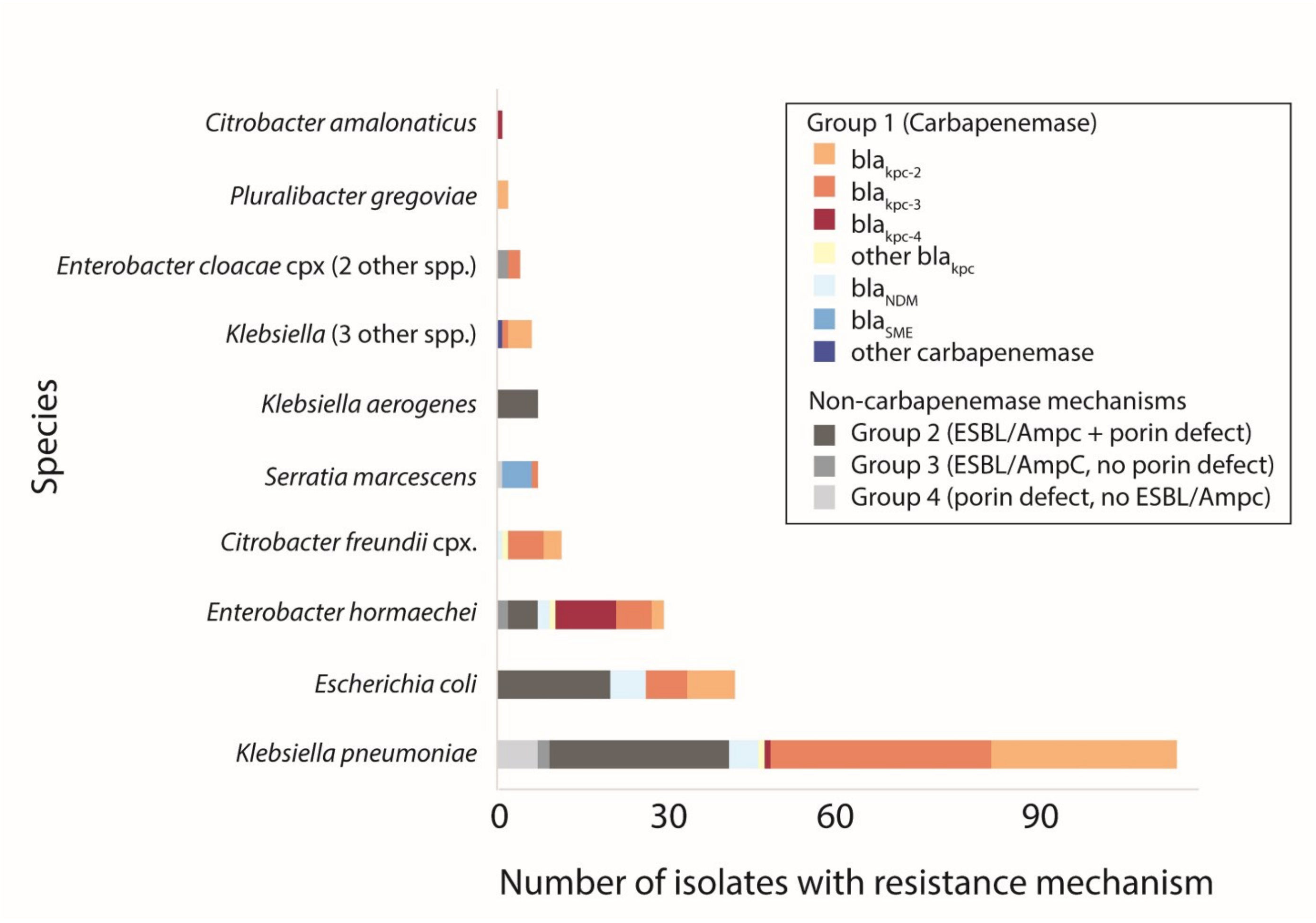


**Figure S2. Resistance mechanisms were diverse, with many shared across species.** The numbers of resistant isolates by resistance mechanism (different colors) are depicted by species.

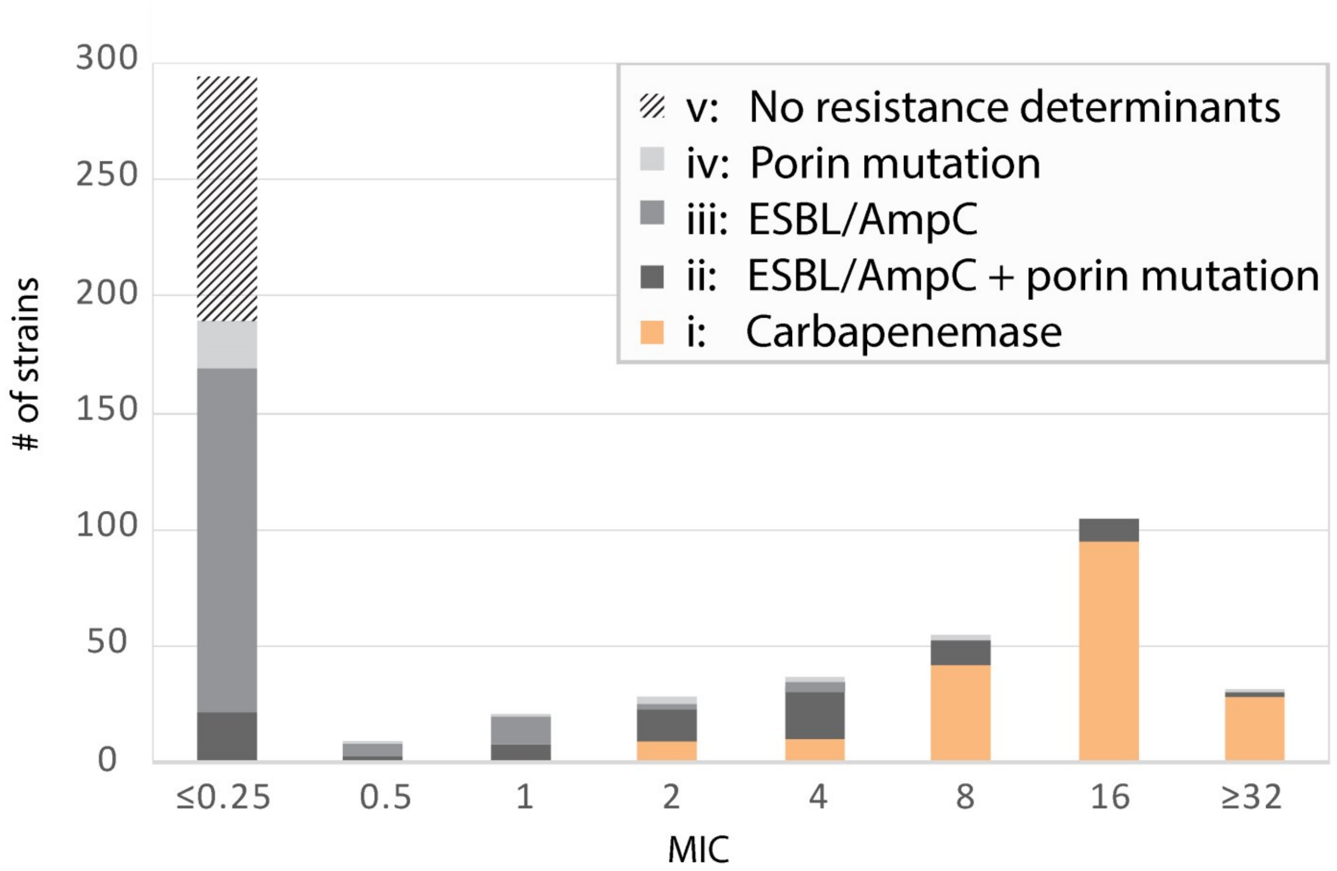


**Figure S3. Carbapenemase-carrying isolates tended to have a higher minimum inhibitory concentration than those with other resistance mechanisms.** Minimum inhibitory concentrations (MICs) of the isolates in our collection are depicted, stratified by mechanism of resistance (different colors).

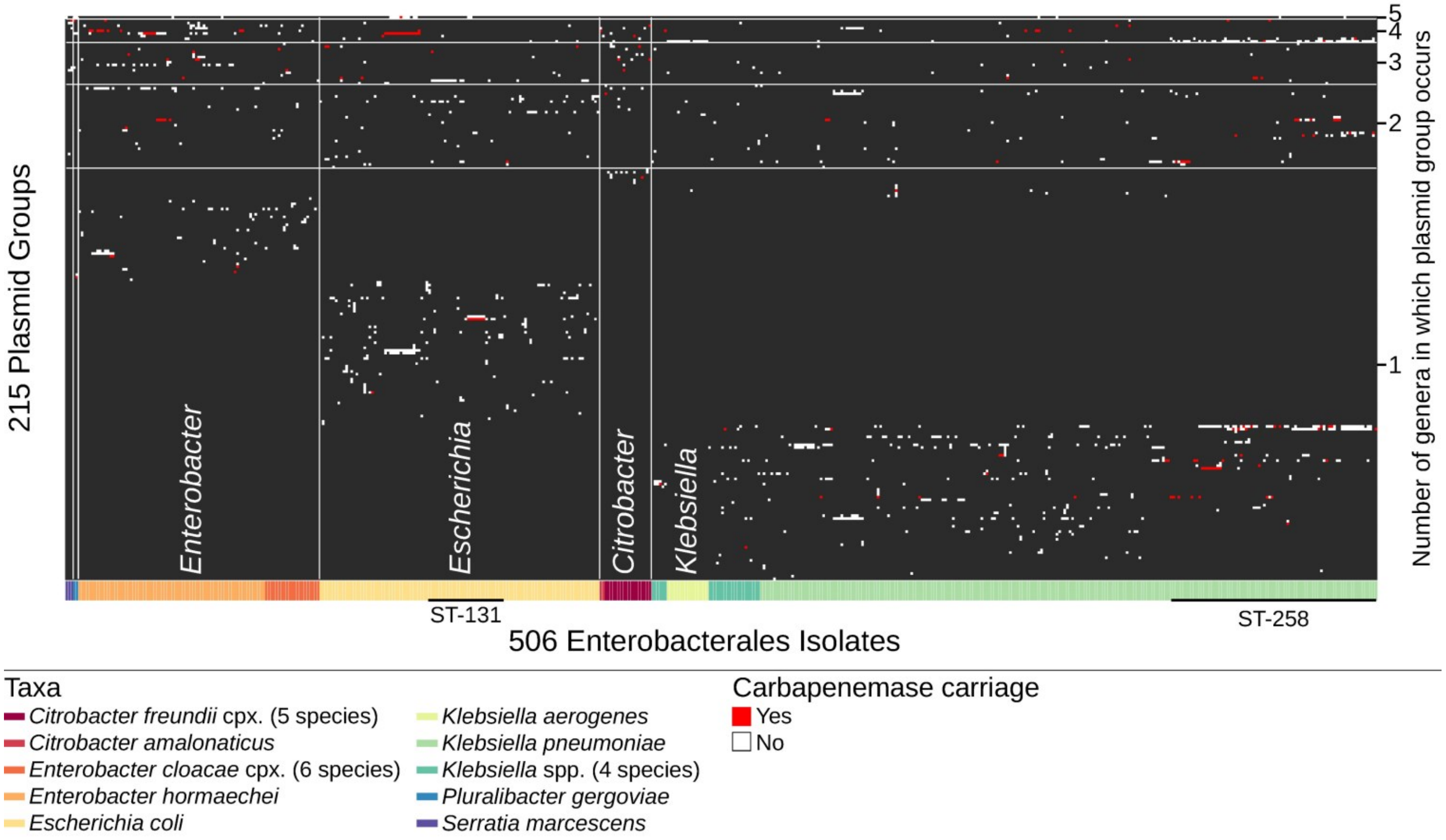


**Figure S4. High level of diversity and phylogenetic range among predicted plasmids.** This plot displays the 215 plasmid groups (rows) contained in all 506 isolates (columns) for which plasmids were predicted. Plasmids with carbapenemases are indicated in red, and plasmids without carbapenemases are indicated in white. Isolates are ordered phylogenetically, while the plasmid groups are ordered by the number of genera in which they occurred and clustered.

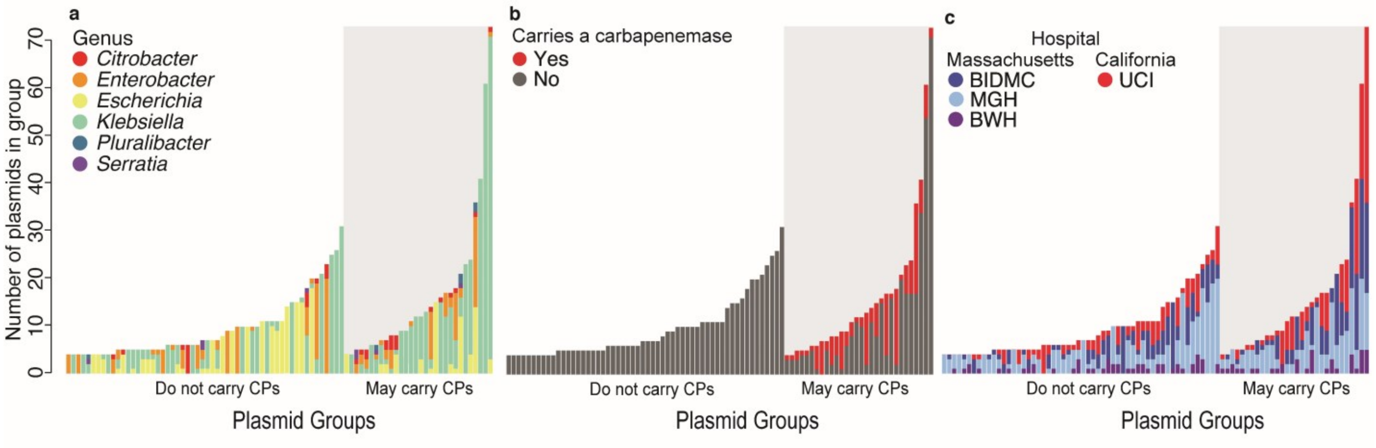


**Figure S5: Plasmids of diverse groups carried carbapenemases and were found in different species, and hospitals.** The number of plasmids from groups for which we found at least four instances is shown. Groups with plasmids that carry carbapenemases (CPs) are depicted on a grey background, while those not observed to carry carbapenemases are shown on a white background. **a**, Plasmid instances colored by carbapenemase carriage. **b**, Plasmid instances colored by genus. **c**, Plasmid groups colored by hospital of origin.

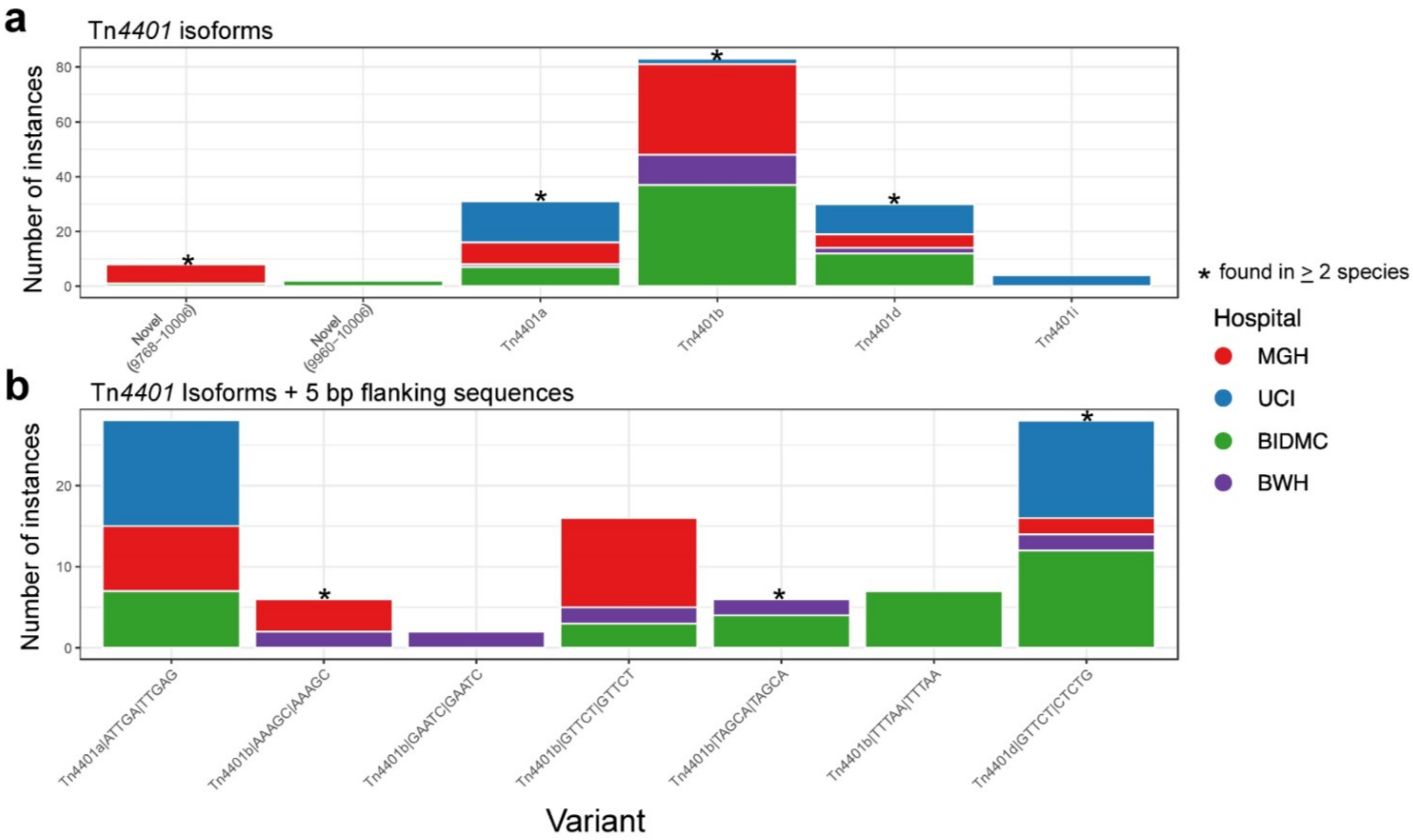


**Figure S6: Limited tracing of carbapenemase localized spread using Tn*4401* isoforms and their immediate flanking sequences. a**, Number of instances for Tn*4401* isoforms; and **b**, combinations of the three most common Tn*4401* isoforms with their 5 bp flanking sequences which were found in multiple isolates from our study. Colors indicate the proportion of instances found in each of the four hospitals. The asterisk indicates forms that were found in multiple species.

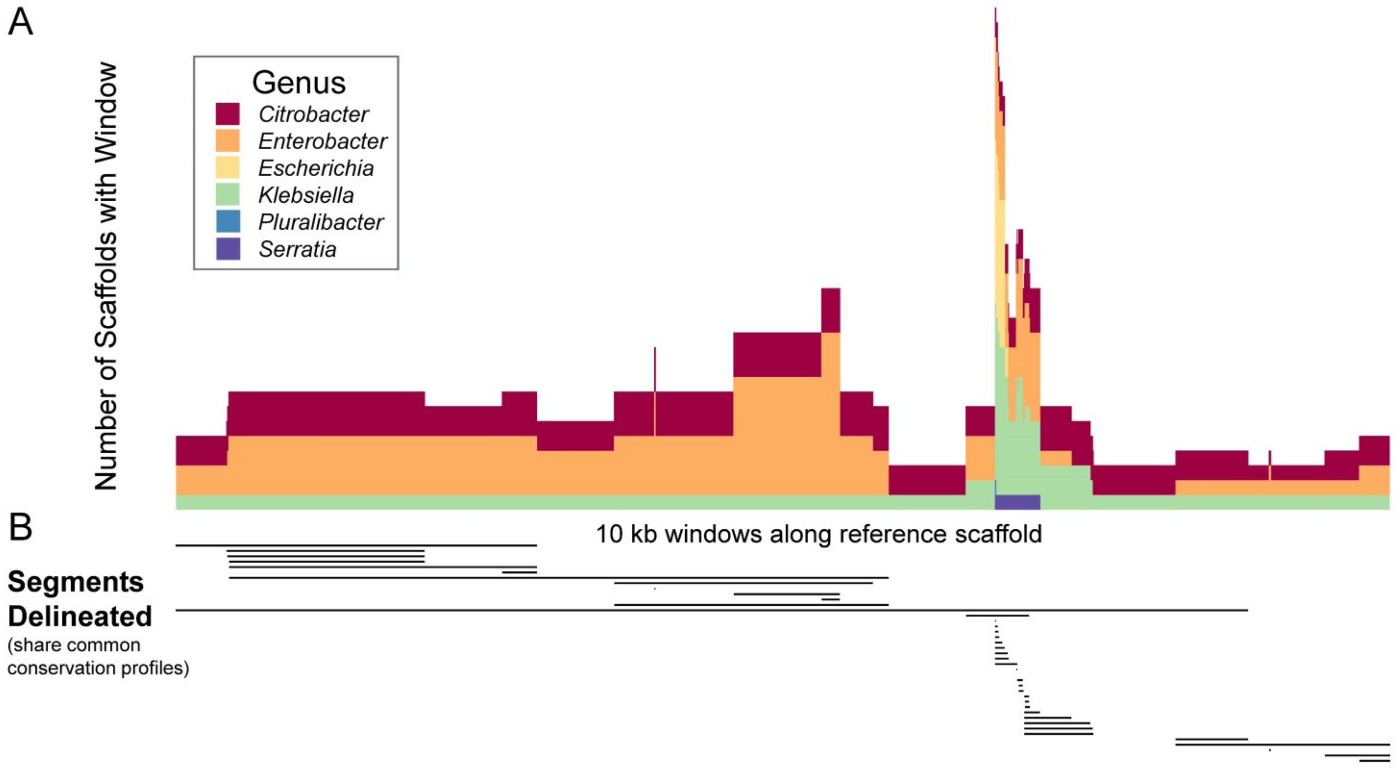


**Figure S7: Method for the delineation of segments shared between plasmids.** *ConSequences* identifies the boundaries of conserved segments spanning multiple 10 kb windows which can be found across multiple (> 2) isolates through assessment of conservation profiles across adjacent windows along reference scaffolds. **a**, Each bar depicts a 10 kb window highlighted by sliding window analysis as being conserved in multiple scaffolds. These bars are ordered along the reference scaffold positionally (x-axis) and the height of bars corresponds to the number of scaffolds in our isolate assemblies that have a highly similar match (≥ 99%) to the 10 kb sequence on the reference (colored by genus). **b**, Using a custom algorithm (Materials and Methods), segments ≥ 10 kb were delineated along the reference scaffold based on conservation profiles across multiple adjacent windows.

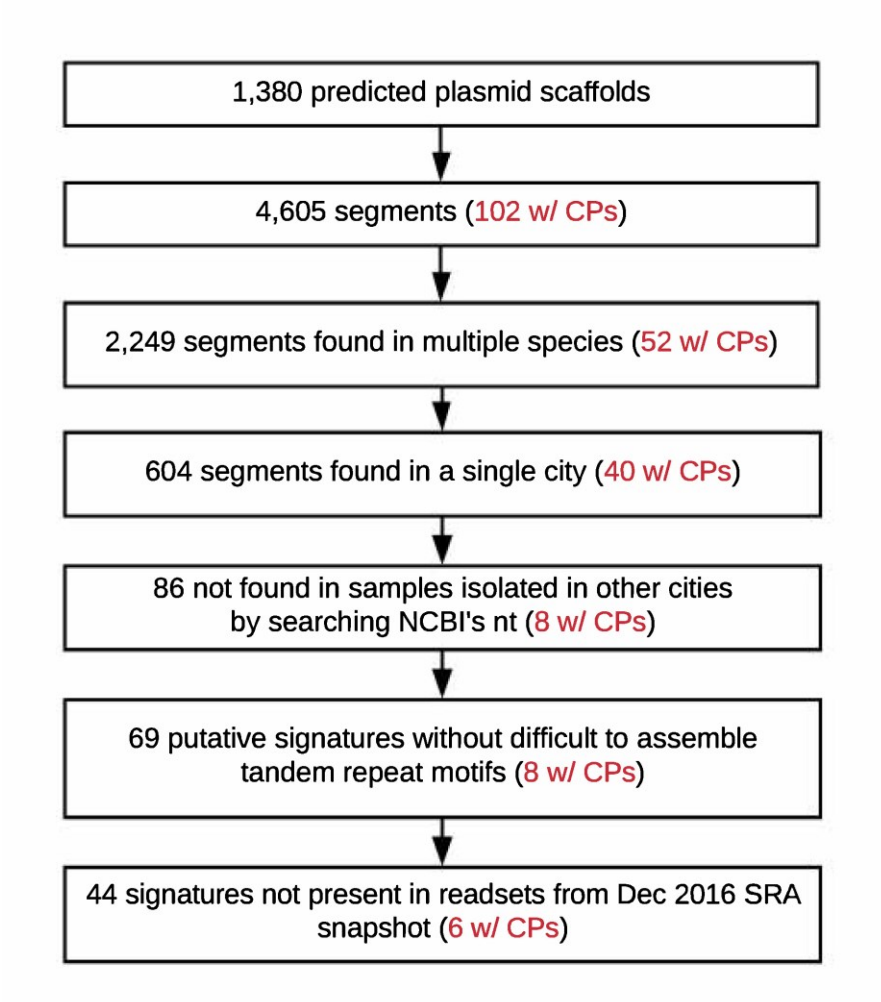


**Figure S8: Workflow to identify geographic signatures.** The number of plasmid segments that were retained after sequentially applying different filters to identify 44 geographic signatures is shown (Materials and Methods). The number of plasmid segments carrying carbapenemases (CPs) is provided in red.

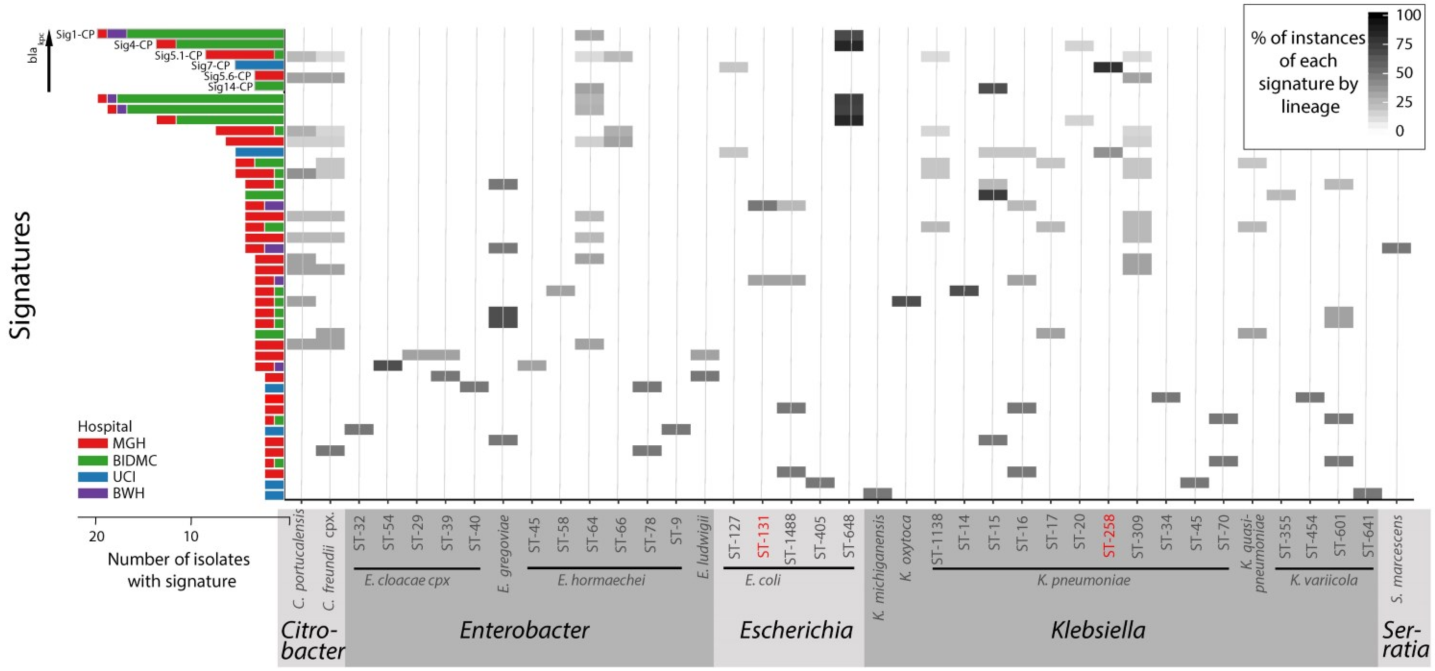


**Figure S9. Signatures were present across diverse sequence types.** In this heatmap, each row corresponds to a unique signature, and each column corresponds to a sequence type (ST). The shading represents the percentage of signature instances belonging to different taxonomic lineages, species or ST. The bar plot to the left of the heatmap depicts the number of isolates containing each signature, highlighting their prevalence across different hospitals.

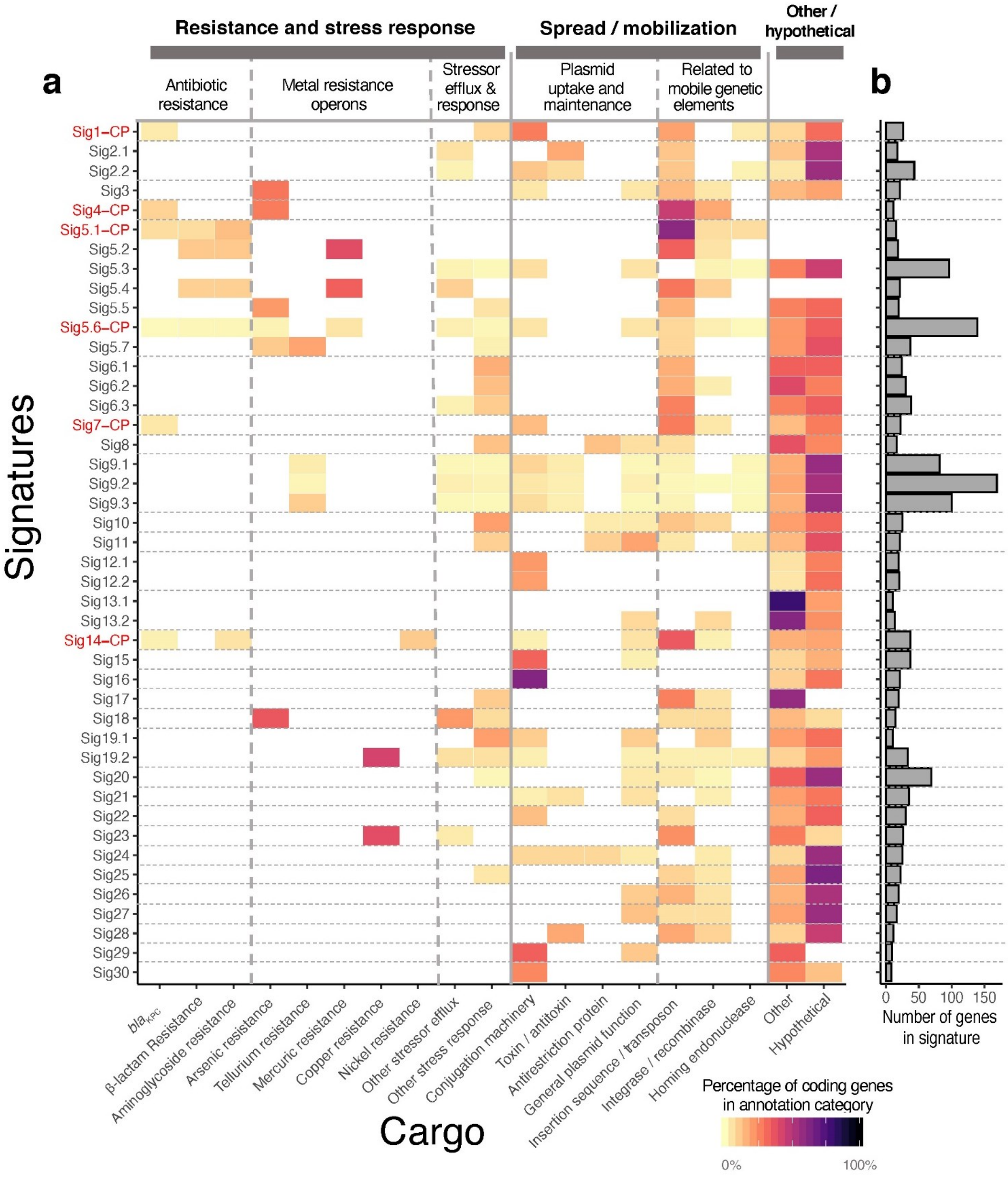


**Figure S10. Signatures carried genes important for hospital adaptation and signature mobility. a**, Each row in the heatmap corresponds to one of the 44 geographic signatures. Groups of signatures that nest into each other are separated by horizontal dashed lines. The predicted functions of 1,494 genes within our 44 signatures were categorized into five major functional categories, unless they fell outside of these categories (other) or no gene function could be predicted (hypothetical). The coloring of the heatmap indicates the percentage of genes of each signature that are assigned to a particular category. The identifiers of carbapenemase-carrying signatures are shown in red type and suffixed with *-CP*. **b**. Number of genes in each signature.

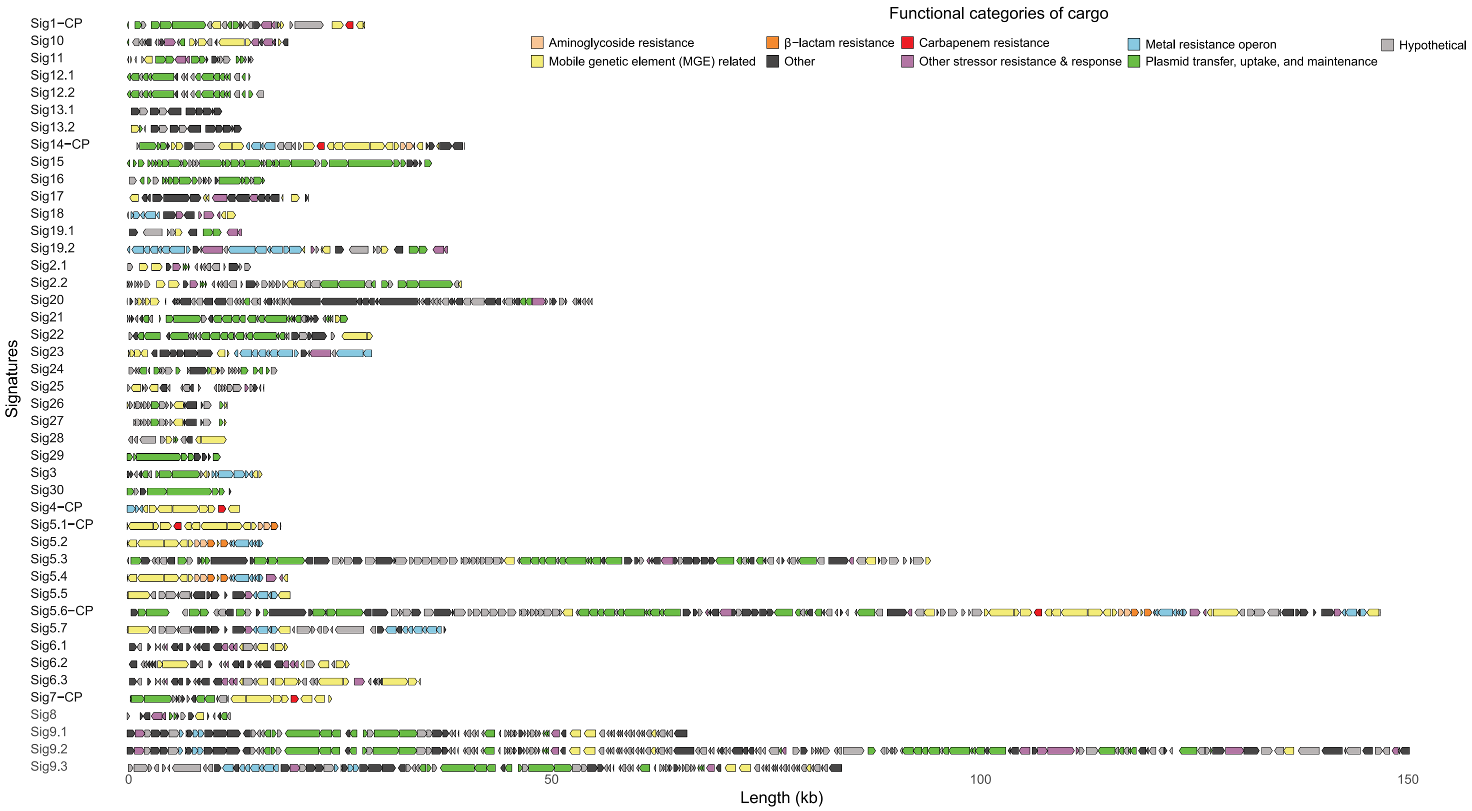


**Figure S11: Details of signature content.** Schematics are shown for the gene content of each signature, including the five with *bla*KPC. Genes are colored according to broad functional categorizations (Materials and Methods).

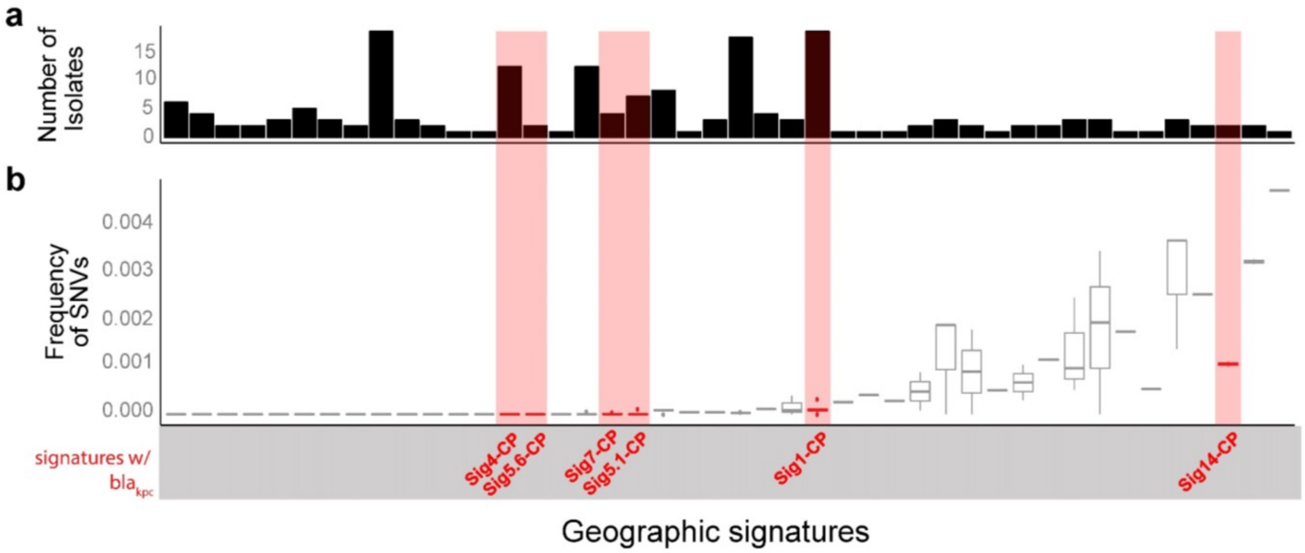


**Figure S12: Signatures were highly conserved and likely derived from a common ancestral sequence.** The number of confident, unambiguous single nucleotide variants (SNVs) differentiating signature instances was calculated for each of the 44 geographic signatures, through comparison of each instance to the signature's representative sequence using Pilon [110]. a, Number of isolates carrying each signature. b, Box plot of SNV frequencies. SNV frequencies were calculated by normalizing the count of SNVs between each signature instance and the reference sequence (c) with the signature's length.

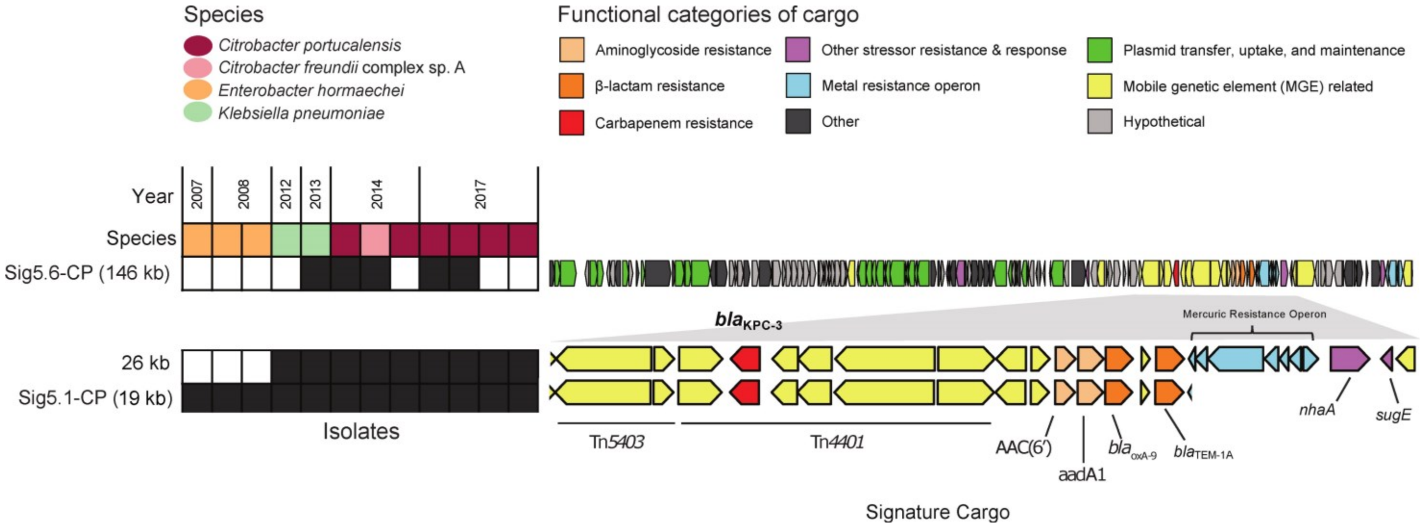


**Figure S13. Geographic signatures with *bla*KPC can occur in multiple configurations across several species and plasmid groups.** The heatmap on the left indicates the presence of signatures Sig5.6-CP and Sig5.1-CP, and the alternate boundaries of the latter, across the twelve isolates found to harbor the signature(s). The gene content of each signature is shown on the right.

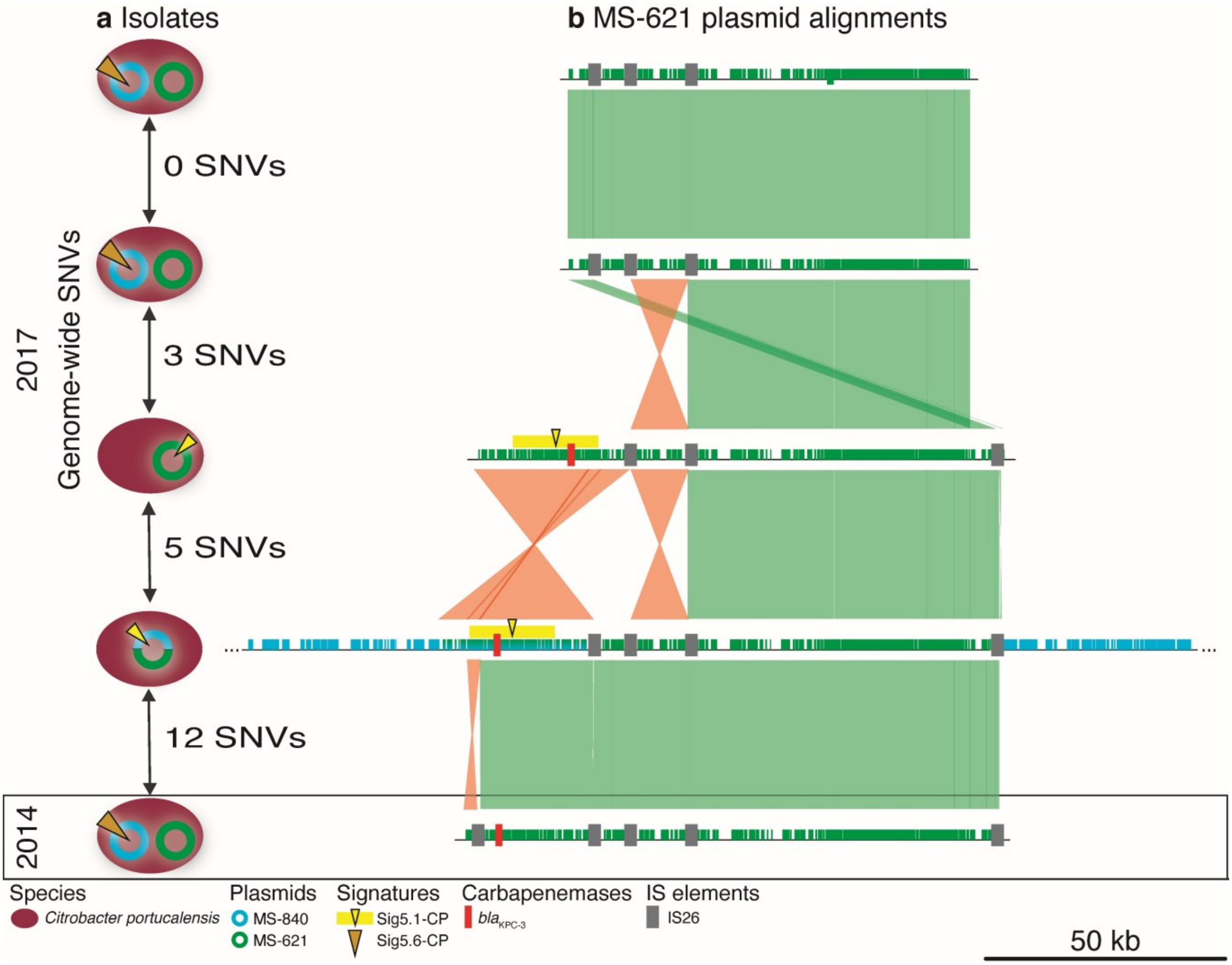


**Figure S14. Geographic signatures with *bla*$_{\mathbf{KPC}}$ occurred in multiple configurations across several species and plasmid groups. a,** The core genome single-nucleotide variants (SNVs) and plasmid and geographic signature carriage of five nearly identical *Citrobacter portucalensis* isolates is shown. **b,** Alignment of the MS-621 plasmids carried by all isolates **a**. Two of these plasmids carry Sig5.1-CP, indicated with the bright yellow bars and triangles. The locations of the *bla*$_{KPC}$ and of insertion sequence IS26 are indicated with red and grey rectangles, respectively. Inversions in the alignment are indicated with orange connector lines; matching regions are indicated with green connector lines. In one isolate, plasmids MS-840 and MS-621 cointegrated, which is indicated by blue alignment flanks.
