## Supplementary results for "Inter-species geographic signatures for tracing horizontal gene transfer and long-term persistence of carbapenem resistance"

**Carbapenemases, as well as extended spectrum beta-lactamases coinciding with disrupted major porins, accounted for the vast majority of carbapenem resistance**

Given that we captured all CRE from hospital microbiology labs, regardless of resistance mechanism, we sought to quantify the contributions of different resistance mechanisms to CRE burden (Figure 2; Figure S1-S2; Table 2; Table S1). To do this, we searched all of our whole genome assemblies, including CRE and CSE, against a database of known genetic determinants of carbapenem resistance, including carbapenemases and extended spectrum beta lactamases (ESBLs), and evaluated the major porins OmpC and OmpF (or their homologues OmpK36 and OmpK35 in *Klebsiella*) for genetic evidence of inactivation [1–3] (Materials and Methods). Based on our findings, we classified each CRE and CSE isolate into one of five previously defined resistome groups [4] (Table 2).

Most CRE (65.9%) carried a carbapenemase (carbapenemase producing CRE or CP-CRE; group i), similar to what was observed in other U.S. and international studies [4,5]; *bla*$_{\mathrm{KPC}}$ variants were among the most prevalent carbapenemases within this group. The small number of group i strains that represented CSE (2%) had MIC's of 0.5 or 1, below but close to our threshold for defining resistance, 2µg/ml. An additional 27.7% of CRE isolates carried a gene encoding an ESBL or AmpC beta-lactamase (Table S5), together with a mutation(s) predicted to inactivate at least one major porin [1–3] (Tables S6-S7) (group ii). About a third of group ii strains were CSE, which may reflect the lack of ESBL upregulation, a known requirement for carbapenem resistance, in these isolates [6–8]. The majority of these strains had very low MIC's (≤0.25). Consistent with the observation that ESBL-producing *Enterobacterales* have increased over time in U.S. hospitals [9], we also observed a significant increase ($p=0.003$; regression slope test) over time in the proportion of CRE carrying *bla*$_{\mathrm{CTX\text{-}M\text{-}15}}$, the dominant ESBL leading to carbapenem resistance in our dataset, and the dominant ESBL worldwide [10] (2.7-fold more CRE carrying *bla*$_{\mathrm{CTX\text{-}M\text{-}15}}$ in 2016 than in 2012; Table S1). We observed a similar statistically significant increase ($p=0.001$) in *bla*$_{CTX-M-15}$ when considering all strains. However, we did not observe this increase when considering the overall proportion of all predicted ESBLs.

As expected, few CRE isolates (2.7%) carried a gene encoding an ESBL or AmpC beta-lactamase in the absence of an inactivating porin mutation (group iii), or had at least one predicted disrupted porin, but no ESBL, AmpC beta-lactamase, or carbapenemase gene (3.6%; group iv). For one of the strains in group iv, BIDMC35, previous work showed that dramatic upregulation of a non-ESBL beta-lactamase (*bla*$_{OXA-663}$) caused resistance, together with a porin disruption [8]. All eight of the CRE isolates in group iv contained at least one non-ESBL beta-lactamase. Additionally, five of these eight isolates were double-porin mutants, which is an established genotype for low-level carbapenem resistance in *Klebsiella* and *Enterobacter* [1]. Taken together, non-CP-CRE (groups ii, iii and iv) had lower median meropenem MIC than CP-CRE (group i) (4 vs. 16; Figure S3), in agreement with previous results [11]. None of the CRE from our dataset fell into group v, representing resistant isolates lacking any of the features captured by groups i-iv. This was fewer than another recent large-scale study of resistance [11].

Although the number of isolates encoding an ESBL or AmpC, but no carbapenemase (groups ii and iii), was similar to that observed in a recent large-scale study of CRE across Europe [4], we observed about twice as many isolates with the corresponding porin disruption necessary to confer resistance (group ii). This was likely due to our intensive search for evidence for genetic inactivation of porins, which was enabled by our sequencing strategy that provided highly contiguous views of the regions around porins, including disruptions caused by transposon insertion. Alternatively, it is possible that there are differences in the distribution of CRE resistance mechanisms between Europe and the U.S. hospitals captured in our study.

**Plasmids with carbapenemases have extensive opportunity for transfer of carbapenemases to new plasmid backgrounds and hosts**

MOB-suite plasmid groups encoding carbapenemases tended to both be observed more often, and to cross genus boundaries more frequently (60% vs. 35% for groups with at least two plasmid instances; Supplementary Figure S4, S5 panel a), than groups that did not contain carbapenemases, either because of sampling or due to an enhanced ability to spread. Perhaps contributing to their relatively high frequency among unrelated organisms, carbapenemase-containing plasmids were more likely than non-carbapenemase plasmids to encode a relaxase, an enzyme required by conjugative plasmids to initiate plasmid transfer [12–14] (93% vs. 79%; $p < 10^{-4}$).

Further, we observed that plasmids from 75% of all groups co-occurred with a plasmid from a carbapenemase-containing plasmid group at some point. These co-occurrences provide ample opportunity for inter-molecular rearrangements between diverse plasmids and diversification of the plasmids within which carbapenemases may be found. Over a quarter (27%) of all isolates with plasmids harbored two or more different carbapenemase-containing plasmid groups, which could lead to plasmid collaboration mechanisms that increase spread of these carbapenemase-carrying groups [15–17]. For instance, nearly half (45%) of the $bla_{KPC}$-carrying isolates with an IncHI2 plasmid also carried an IncF plasmid. These two plasmids conjugate optimally at different temperatures, but co-integration of a plasmid from each group would extend the range of temperatures at which the co-integrate could be shared [17].

2. Vergalli J, Bodrenko IV, Masi M, Moynié L, Acosta-Gutiérrez S, Naismith JH, et al. Porins and small-molecule translocation across the outer membrane of Gram-negative bacteria. Nat Rev

Microbiol. 2020;18:164–76.

Medical Soc; 2020;382:1309–19.
