## Supplementary Figure S1 for "Inter-species geographic signatures for tracing horizontal gene transfer and long-term persistence of carbapenem resistance"

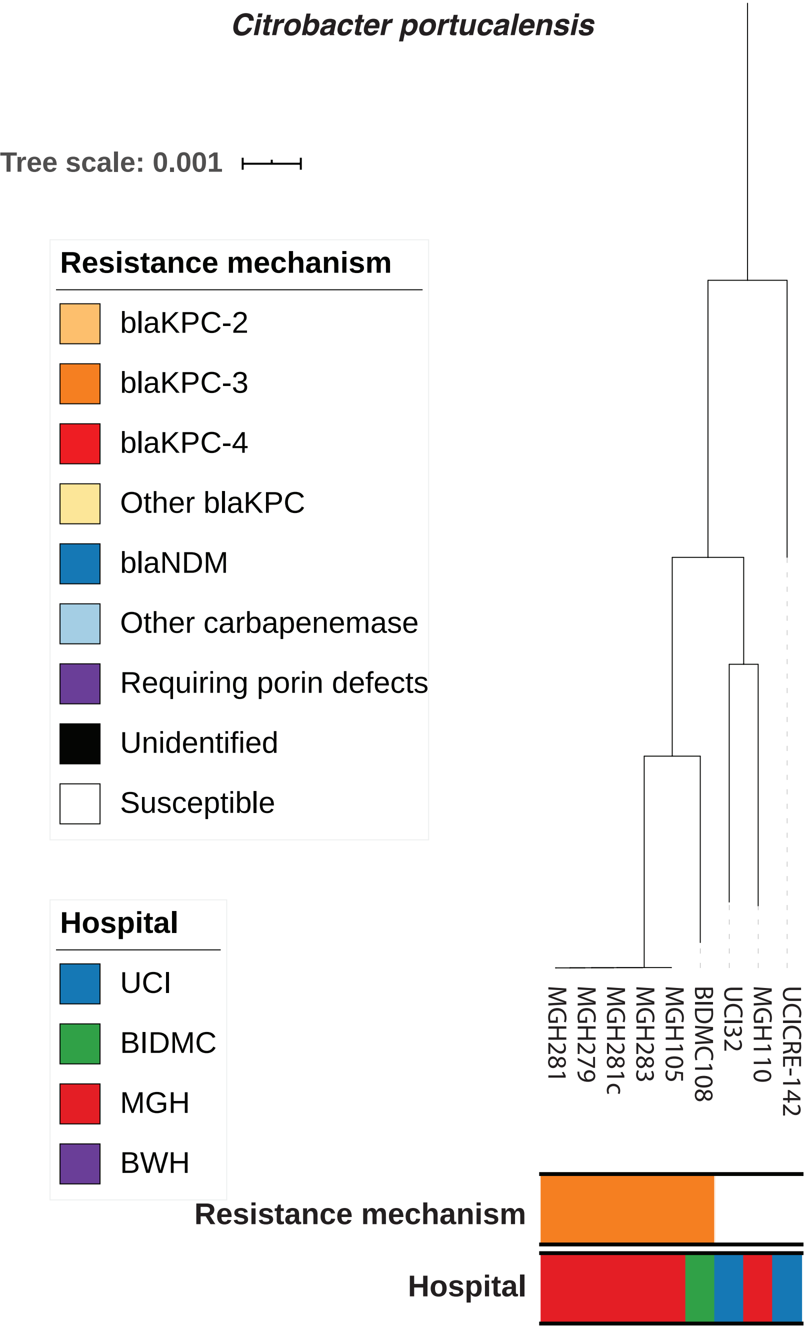
Citrobacter portucalensis
Tree scale: 0.001
Resistance mechanism
blaKPC-2
blaKPC-3
blaKPC-4
Other blaKPC
blaNDM
Other carbapenemase
Requiring porin defects
Unidentified
Susceptible
Hospital
UCI
BIDMC
MGH
BWH
MGH281
MGH279
MGH281c
MGH283
MGH105
BIDMC108
UCI32
MGH110
UCICRE-142
Resistance mechanism
Hospital

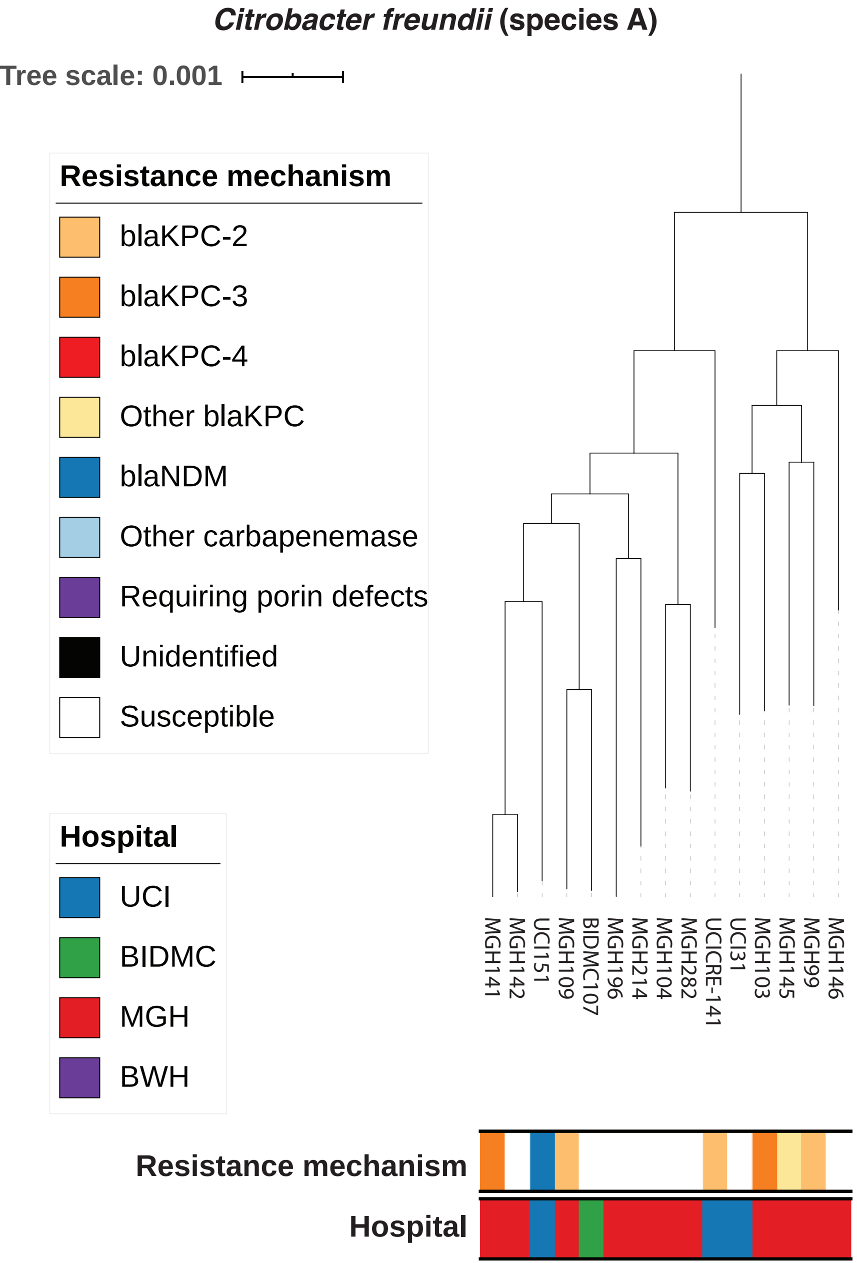
Citrobacter freundii (species A)
Tree scale: 0.001
Resistance mechanism
blaKPC-2
blaKPC-3
blaKPC-4
Other blaKPC
blaNDM
Other carbapenemase
Requiring porin defects
Unidentified
Susceptible
Hospital
UCI
BIDMC
MGH
BWH
MGH141
MGH142
UCI151
MGH109
BIDMC107
MGH196
MGH214
MGH104
MGH282
UCICRE-141
UCI31
MGH103
MGH145
MGH99
MGH146
Resistance mechanism
Hospital

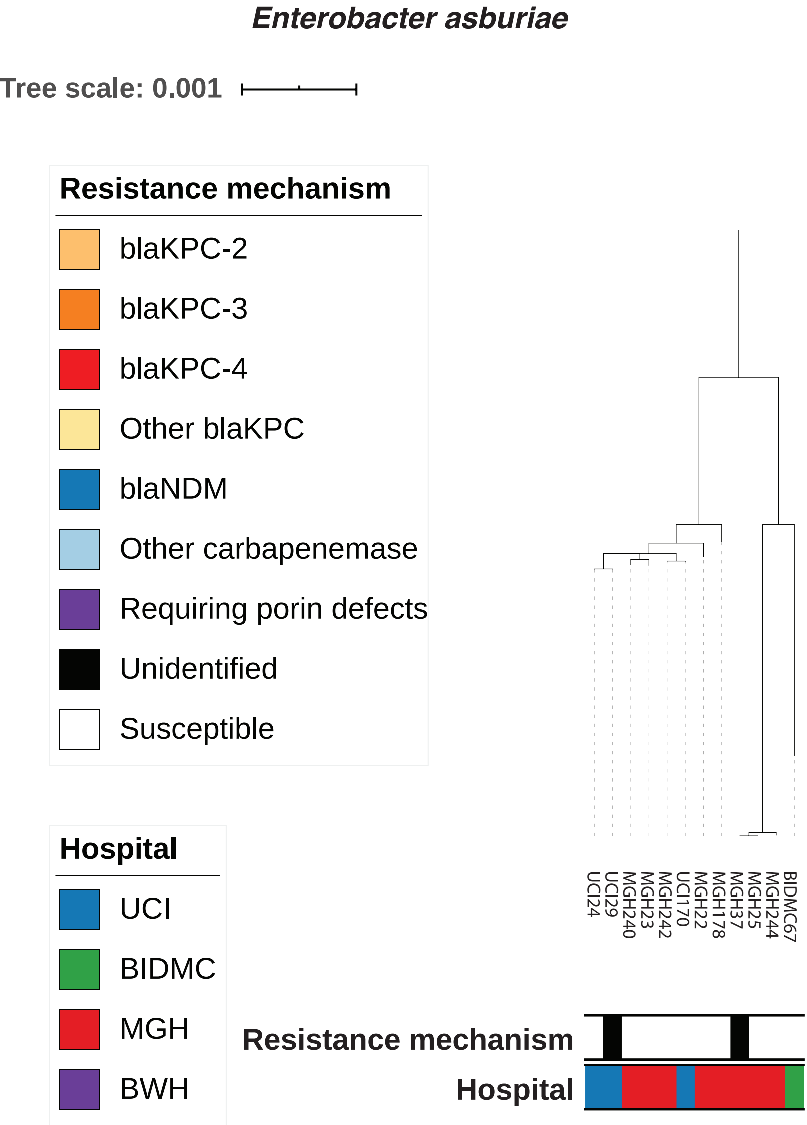
Enterobacter asburiae
Tree scale: 0.001
Resistance mechanism
blaKPC-2
blaKPC-3
blaKPC-4
Other blaKPC
blaNDM
Other carbapenemase
Requiring porin defects
Unidentified
Susceptible
Hospital
UCI
BIDMC
MGH
BWH
UCI24
UCI29
MGH240
MGH23
MGH242
UCI170
MGH22
MGH178
MGH37
MGH25
MGH244
BIDMC67
Resistance mechanism
Hospital

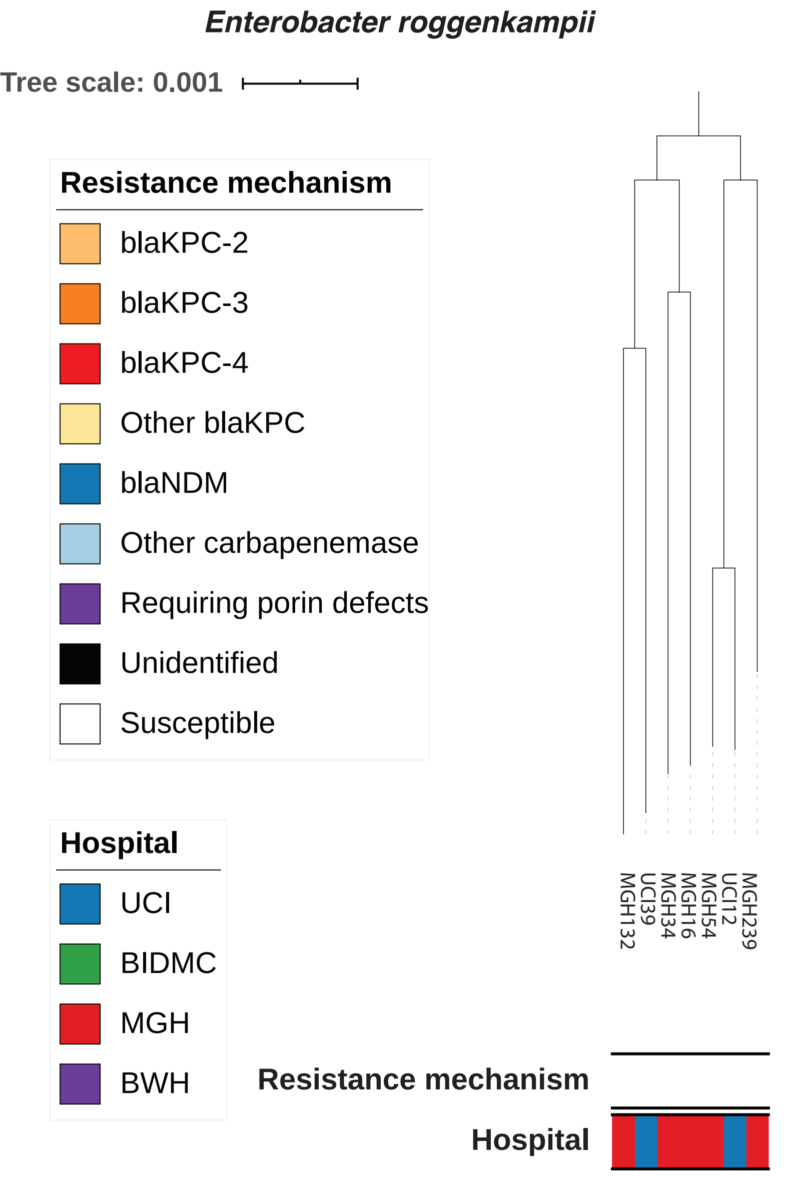
Enterobacter roggenkampii
Tree scale: 0.001
Resistance mechanism
blaKPC-2
blaKPC-3
blaKPC-4
Other blaKPC
blaNDM
Other carbapenemase
Requiring porin defects
Unidentified
Susceptible
Hospital
UCI
BIDMC
MGH
BWH
MGH132
UCI39
MGH34
MGH16
MGH54
UCI12
MGH239
Resistance mechanism
Hospital

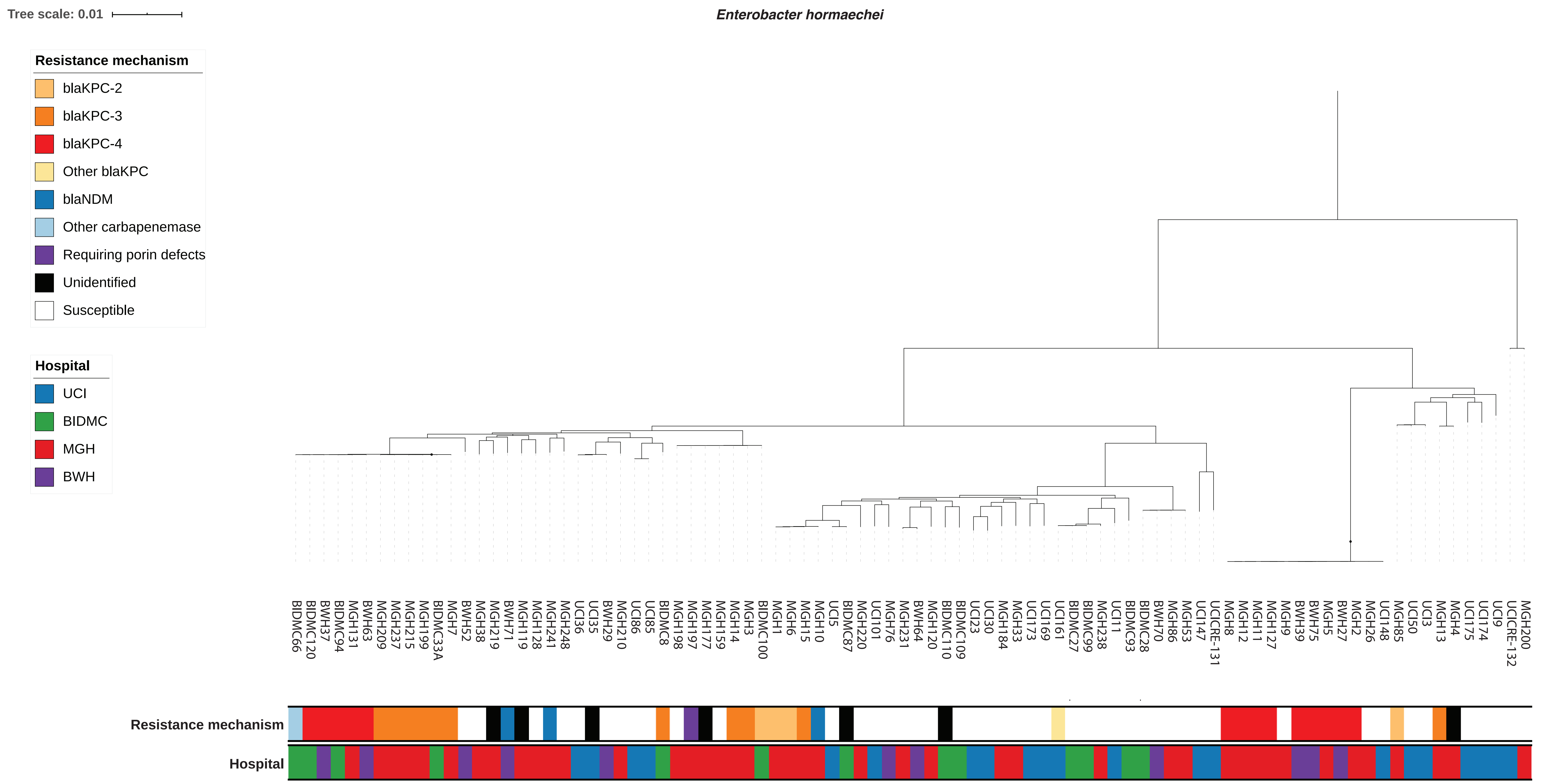
Tree scale: 0.01
Enterobacter hormaechei
Resistance mechanism
blaKPC-2
blaKPC-3
blaKPC-4
Other blaKPC
blaNDM
Other carbapenemase
Requiring porin defects
Unidentified
Susceptible
Hospital
UCI
BIDMC
MGH
BWH
BIDMC66 BIDMC120 BWH37 BIDMC94 MGH131 BWH63 MGH209 MGH237 MGH215 MGH199 BIDMC33A MGH7 BWH52 MGH38 MGH219 BWH71 MGH119 MGH128 MGH241 MGH248 UCI36 UCI35 BWH29 MGH210 UCI86 UCI85 BIDMC8 MGH198 MGH197 MGH177 MGH159 MGH14 MGH3 BIDMC100 MGH1 MGH6 MGH15 MGH10 UCI5 BIDMC87 MGH220 UCI101 MGH76 MGH231 BWH64 MGH120 BIDMC110 BIDMC109 UCI23 UCI30 MGH184 MGH33 UCI173 UCI169 UCI161 BIDMC27 BIDMC99 MGH238 UCI11 BIDMC93 BIDMC28 BWH70 MGH86 MGH53 UCI147 UCICRE-131 MGH8 MGH12 MGH11 MGH127 MGH9 BWH39 BWH75 MGH5 BWH27 MGH2 MGH26 UCI148 MGH85 UCI50 UCI3 MGH13 MGH4 UCI175 UCI174 UCI9 UCICRE-132 MGH200
Resistance mechanism
Hospital

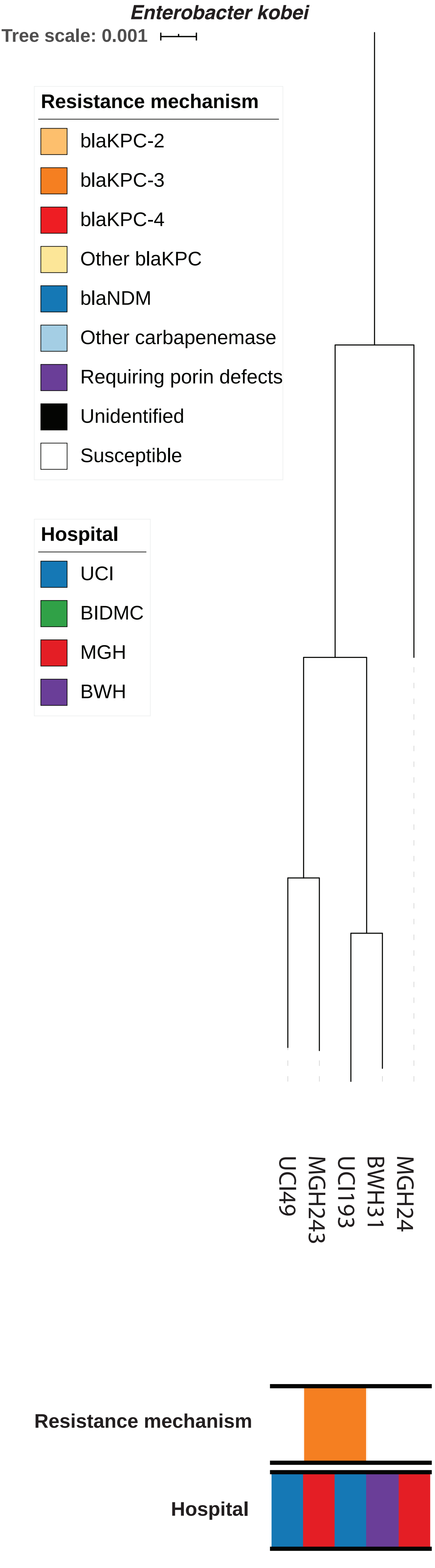
Enterobacter kobei
Tree scale: 0.001
Resistance mechanism
blaKPC-2
blaKPC-3
blaKPC-4
Other blaKPC
blaNDM
Other carbapenemase
Requiring porin defects
Unidentified
Susceptible
Hospital
UCI
BIDMC
MGH
BWH
UCI49
MGH243
UCI193
BWH31
MGH24
Resistance mechanism
Hospital

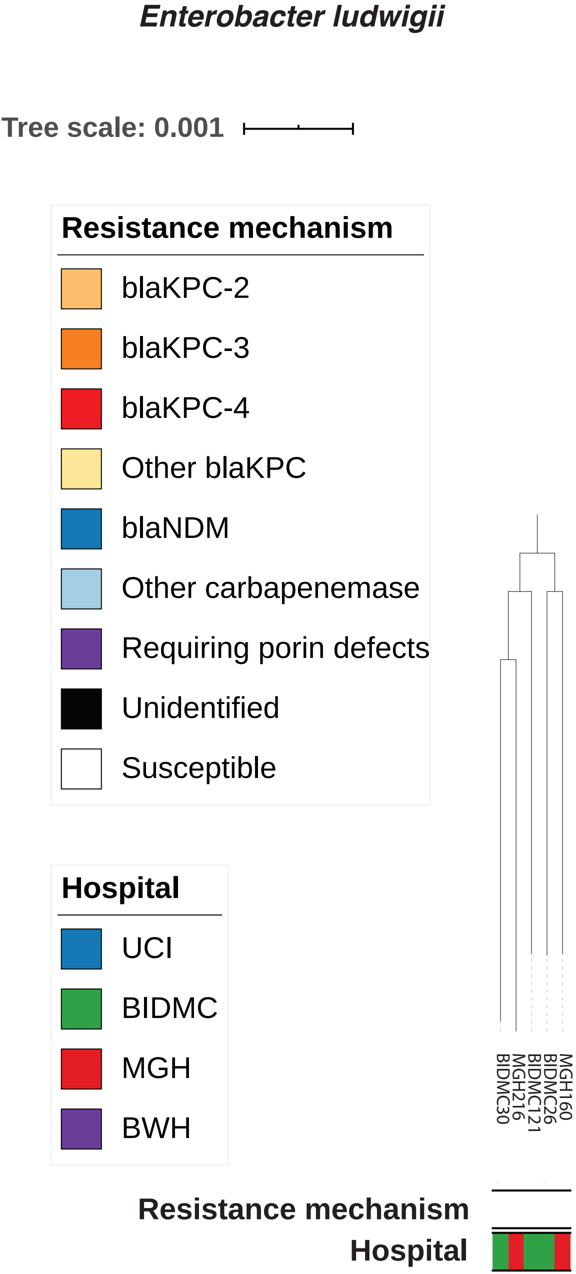
Enterobacter ludwigii
Tree scale: 0.001
Resistance mechanism
blaKPC-2
blaKPC-3
blaKPC-4
Other blaKPC
blaNDM
Other carbapenemase
Requiring porin defects
Unidentified
Susceptible
Hospital
UCI
BIDMC
MGH
BWH
BIDMC30
MGH216
BIDMC121
BIDMC26
MGH160
Resistance mechanism
Hospital

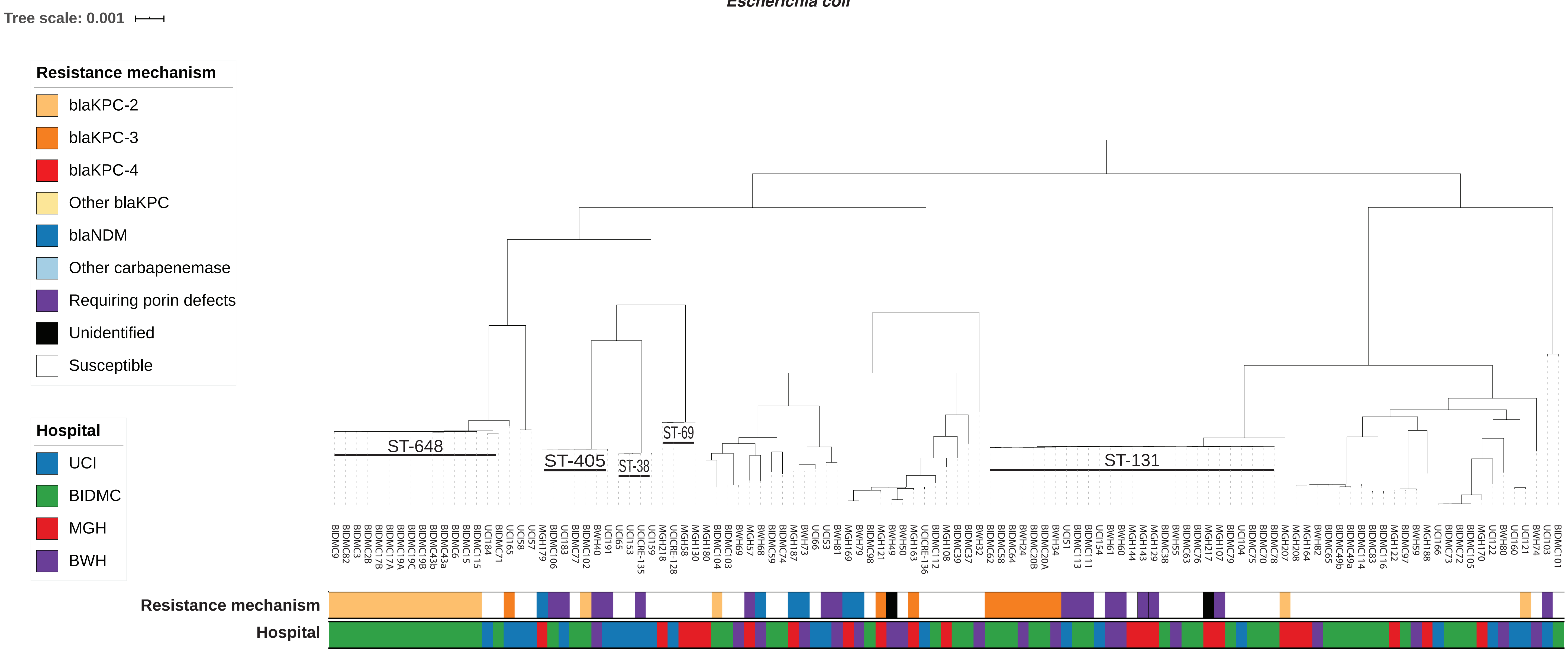

Escherichia coli
Tree scale: 0.001
Resistance mechanism
blaKPC-2
blaKPC-3
blaKPC-4
Other blaKPC
blaNDM
Other carbapenemase
Requiring porin defects
Unidentified
Susceptible
Hospital
UCI
BIDMC
MGH
BWH
ST-648
ST-405
ST-38
ST-69
ST-131
Resistance mechanism
Hospital

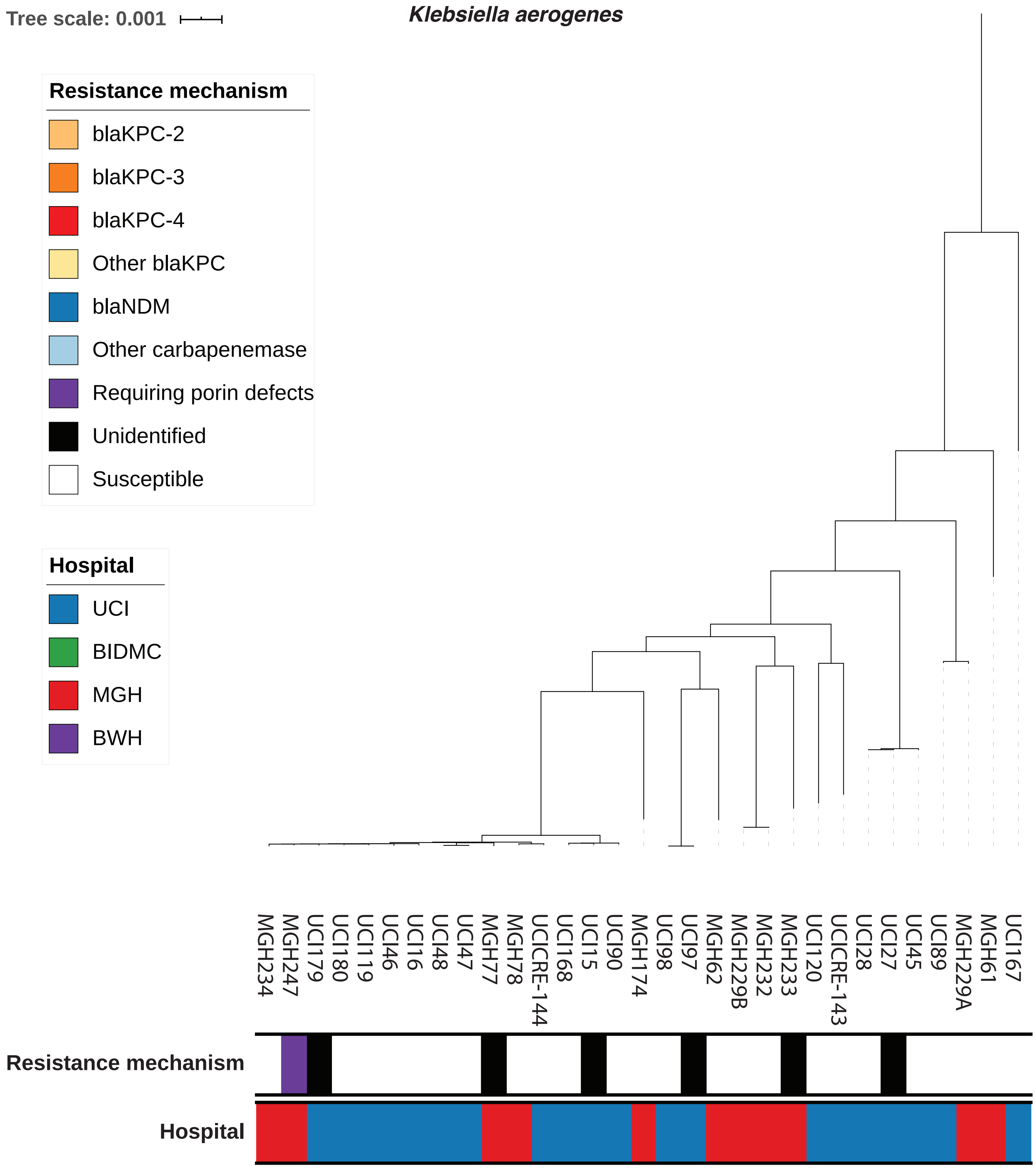
Tree scale: 0.001
Klebsiella aerogenes
Resistance mechanism
blaKPC-2
blaKPC-3
blaKPC-4
Other blaKPC
blaNDM
Other carbapenemase
Requiring porin defects
Unidentified
Susceptible
Hospital
UCI
BIDMC
MGH
BWH
MGH234
MGH247
UCI179
UCI180
UCI119
UCI46
UCI16
UCI48
UCI47
MGH77
MGH78
UCICRE-144
UCI168
UCI15
UCI90
MGH174
UCI98
UCI97
MGH62
MGH229B
MGH232
MGH233
UCI120
UCICRE-143
UCI28
UCI27
UCI45
UCI89
MGH229A
MGH61
UCI167
Resistance mechanism
Hospital

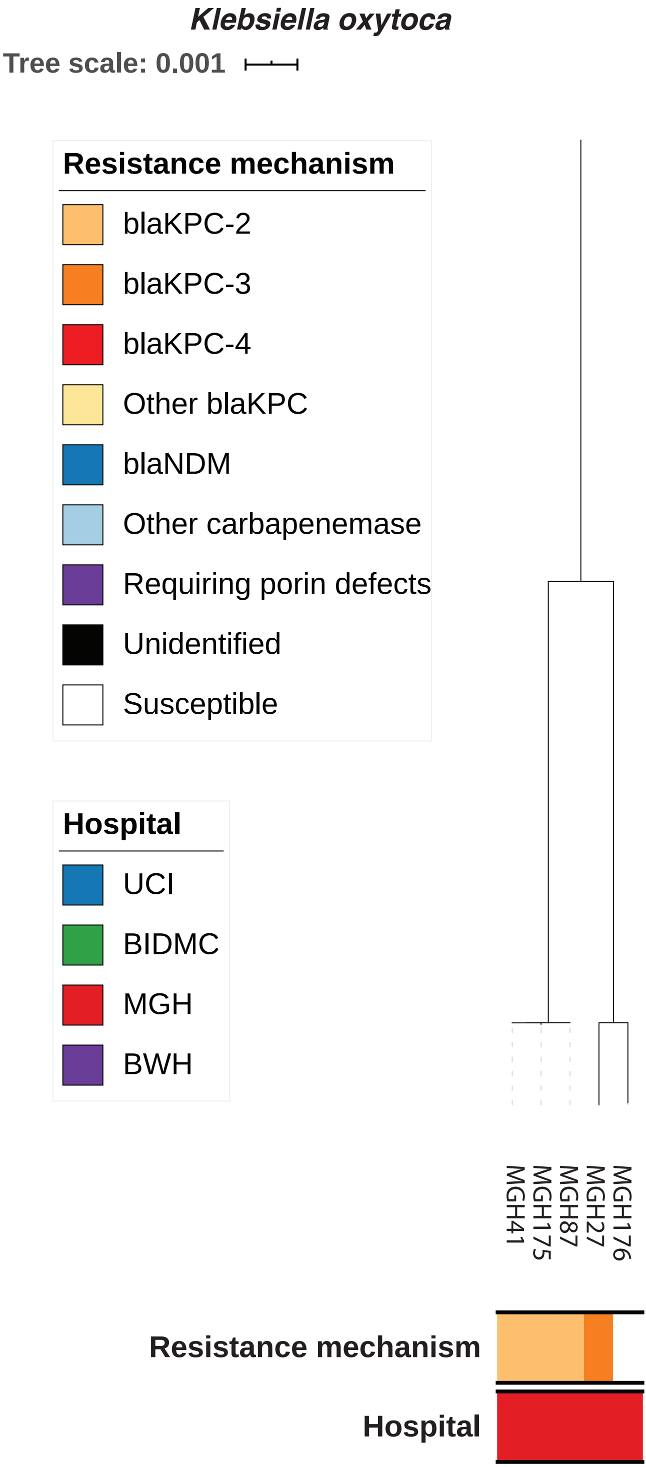
Klebsiella oxytoca
Tree scale: 0.001
Resistance mechanism
blaKPC-2
blaKPC-3
blaKPC-4
Other blaKPC
blaNDM
Other carbapenemase
Requiring porin defects
Unidentified
Susceptible
Hospital
UCI
BIDMC
MGH
BWH
MGH41
MGH175
MGH87
MGH27
MGH176
Resistance mechanism
Hospital

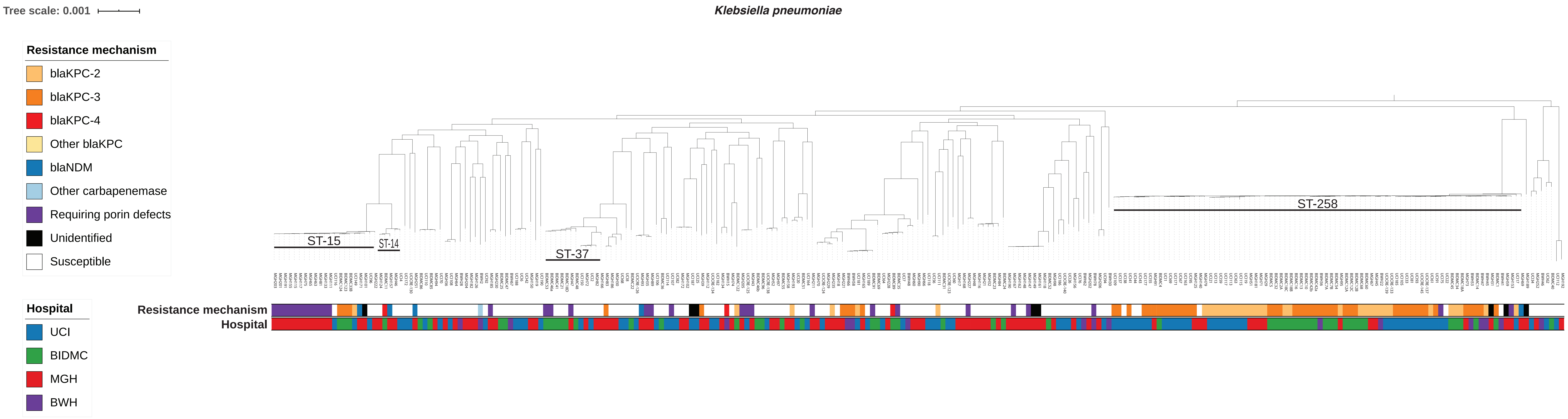

Tree scale: 0.001
Klebsiella pneumoniae
Resistance mechanism
blaKPC-2
blaKPC-3
blaKPC-4
Other blaKPC
blaNDM
Other carbapenemase
Requiring porin defects
Unidentified
Susceptible
ST-15
ST-14
ST-37
ST-258
Resistance mechanism
Hospital
Hospital
UCI
BIDMC
MGH
BWH

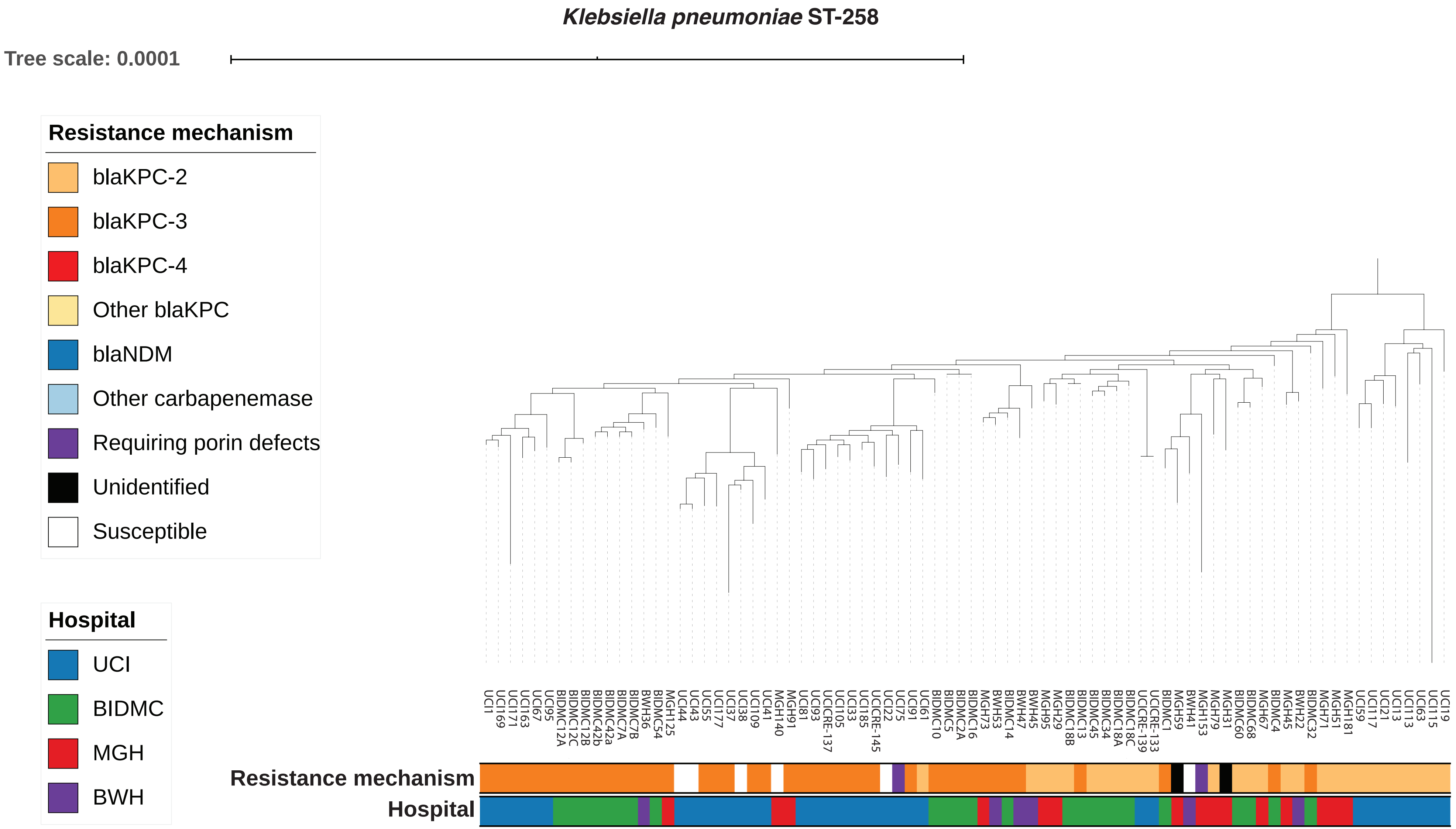
*Klebsiella pneumoniae* ST-258
Tree scale: 0.0001
Resistance mechanism
blaKPC-2
blaKPC-3
blaKPC-4
Other blaKPC
blaNDM
Other carbapenemase
Requiring porin defects
Unidentified
Susceptible
Hospital
UCI
BIDMC
MGH
BWH
UCI1 UCI169 UCI171 UCI163 UCI67 UCI95 BIDMC12A BIDMC12C BIDMC12B BIDMC42b BIDMC42a BIDMC7A BIDMC7B BWH36 BIDMC54 MGH125 UCI44 UCI43 UCI55 UCI177 UCI37 UCI38 UCI109 UCI41 MGH140 MGH91 UCI81 UCI93 UCICRE-137 UCI105 UCI33 UCI185 UCICRE-145 UCI22 UCI75 UCI91 UCI61 BIDMC10 BIDMC5 BIDMC2A BIDMC16 MGH73 BWH53 BIDMC14 BWH47 BWH45 MGH95 MGH29 BIDMC18B BIDMC13 BIDMC45 BIDMC34 BIDMC18A BIDMC18C UCICRE-139 UCICRE-133 BIDMC1 MGH59 BWH41 MGH153 MGH79 MGH31 BIDMC60 BIDMC68 MGH67 BIDMC4 MGH45 BWH22 BIDMC32 MGH71 MGH51 MGH181 UCI59 UCI117 UCI21 UCI13 UCI113 UCI63 UCI115 UCI19
Resistance mechanism
Hospital

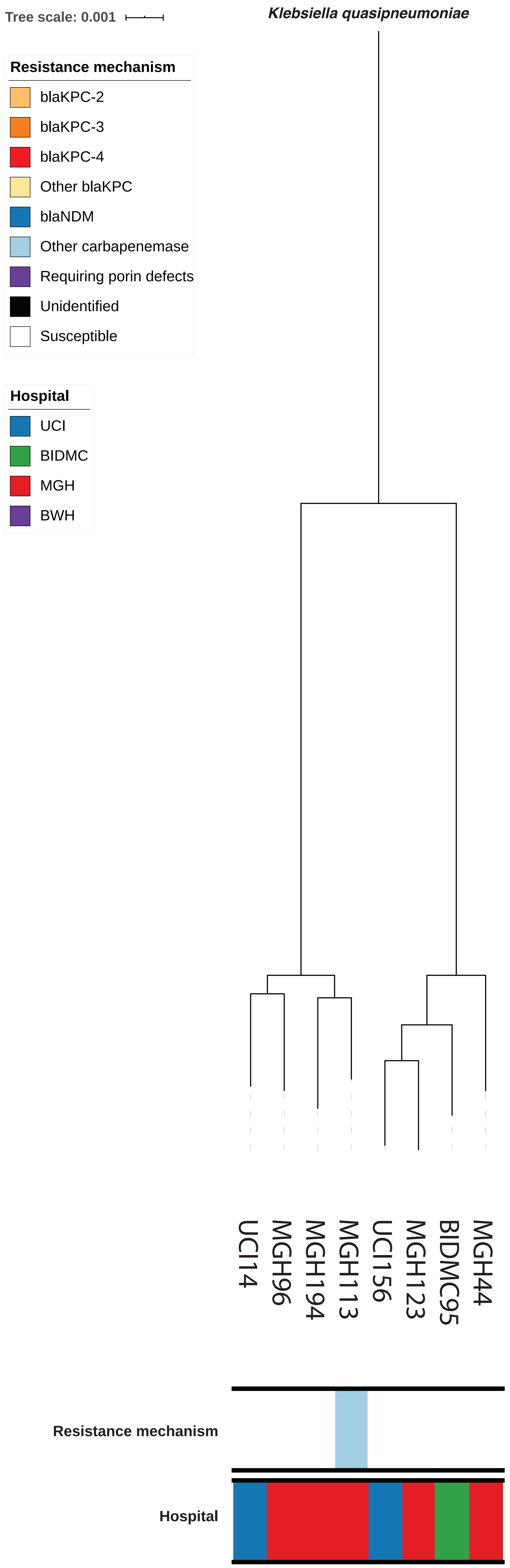

Tree scale: 0.001
Klebsiella quasipneumoniae
Resistance mechanism
blaKPC-2
blaKPC-3
blaKPC-4
Other blaKPC
blaNDM
Other carbapenemase
Requiring porin defects
Unidentified
Susceptible
Hospital
UCI
BIDMC
MGH
BWH
UCI14
MGH96
MGH194
MGH113
UCI156
MGH123
BIDMC95
MGH44
Resistance mechanism
Hospital

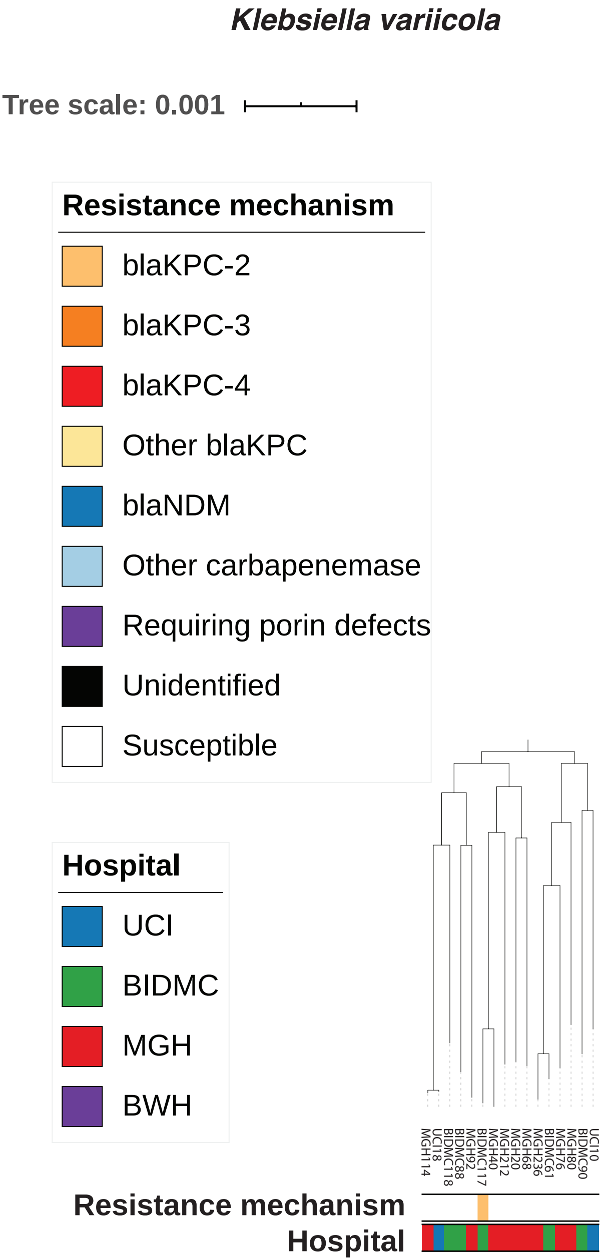

Klebsiella variicola
Tree scale: 0.001
Resistance mechanism
blaKPC-2
blaKPC-3
blaKPC-4
Other blaKPC
blaNDM
Other carbapenemase
Requiring porin defects
Unidentified
Susceptible
Hospital
UCI
BIDMC
MGH
BWH
MGH114
UCI18
BIDMC118
BIDMC88
MGH92
BIDMC117
MGH40
MGH212
MGH20
MGH68
MGH236
BIDMC61
MGH76
MGH80
BIDMC90
UCI10
Resistance mechanism
Hospital

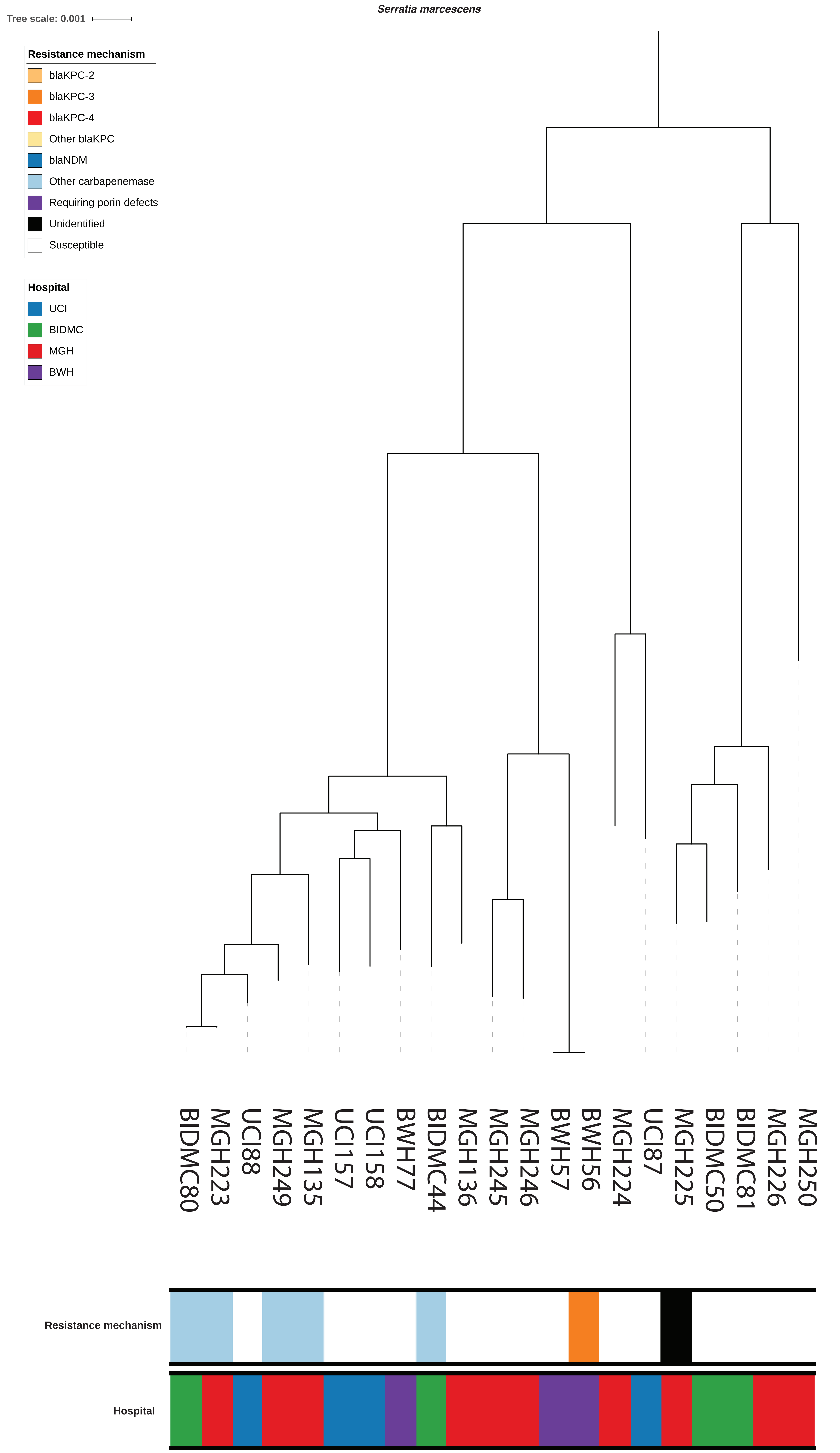
Serratia marcescens
Tree scale: 0.001
Resistance mechanism
blaKPC-2
blaKPC-3
blaKPC-4
Other blaKPC
blaNDM
Other carbapenemase
Requiring porin defects
Unidentified
Susceptible
Hospital
UCI
BIDMC
MGH
BWH
BIDMC80
MGH223
UCI88
MGH249
MGH135
UCI157
UCI158
BWH77
BIDMC44
MGH136
MGH245
MGH246
BWH57
BWH56
MGH224
UCI87
MGH225
BIDMC50
BIDMC81
MGH226
MGH250
Resistance mechanism
Hospital
